# Targeted therapies prime oncogene-driven lung cancers for macrophage-mediated destruction

**DOI:** 10.1101/2023.03.03.531059

**Authors:** Kyle Vaccaro, Juliet Allen, Troy W. Whitfield, Asaf Maoz, Sarah Reeves, José Velarde, Dian Yang, Nicole Phan, George W. Bell, Aaron N. Hata, Kipp Weiskopf

## Abstract

Macrophage immune checkpoint inhibitors, such as anti-CD47 antibodies, show promise in clinical trials for solid and hematologic malignancies. However, the best strategies to use these therapies remain unknown and ongoing studies suggest they may be most effective when used in combination with other anticancer agents. Here, we developed a novel screening platform to identify drugs that render lung cancer cells more vulnerable to macrophage attack, and we identified therapeutic synergy exists between genotype-directed therapies and anti-CD47 antibodies. In validation studies, we found the combination of genotype-directed therapies and CD47 blockade elicited robust phagocytosis and eliminated persister cells in vitro and maximized anti-tumor responses in vivo. Importantly, these findings broadly applied to lung cancers with various RTK/MAPK pathway alterations—including *EGFR* mutations, *ALK* fusions, or *KRAS^G12C^* mutations. We observed downregulation of β2-microglobulin and CD73 as molecular mechanisms contributing to enhanced sensitivity to macrophage attack. Our findings demonstrate that dual inhibition of the RTK/MAPK pathway and the CD47/SIRPa axis is a promising immunotherapeutic strategy. Our study provides strong rationale for testing this therapeutic combination in patients with lung cancers bearing driver mutations.

**Brief summary:** Unbiased drug screens identify targeted therapies as drugs that make lung cancers with driver mutations more vulnerable to macrophage attack.

## INTRODUCTION

Many cancers arise from a single mutation in a proto-oncogene that drives tumor growth.

These cancers are often treated with genotype-directed ‘targeted therapies,’ small molecule inhibitors that specifically disable growth signals from oncogenic driver mutations. These drugs are most frequently used to treat non-small cell lung cancer (NSCLC), where up to 50% of patients may have an actionable mutation that can be targeted with specific small molecule inhibitors (1). *EGFR* and *KRAS* mutations are among the most common actionable mutations in lung cancer (1-4). Although targeted therapies extend patients’ lives, they typically do not cure patients and responses generally last for only a few years (2, 3, 5). For these reasons, there is a critical need to identify more effective ways to treat lung cancers with oncogenic driver mutations.

Based on the success of cancer immunotherapy, an appealing strategy has been to combine targeted therapies with agents that activate the immune system to attack cancer. However, lung cancers with driver mutations may have lower tumor mutational burden and fewer neoantigens, thereby impairing adaptive immune responses (6, 7). Furthermore, the combination of some targeted therapies with anti-PD-1/PD-L1 agents has been limited in efficacy or produced synergistic toxicity (8). In contrast to targeting T cells, macrophages are an attractive alternative target since they are often the most common infiltrating immune cell within the tumor microenvironment (9, 10). Nevertheless, the combination of targeted therapies with macrophage-directed therapies is relatively unexplored.

The CD47/SIRPa interaction is the best-characterized immune checkpoint that controls macrophages in tumors (11). CD47 is highly expressed on many cancers, and it binds to the inhibitory receptor SIRPa that is present on macrophages and other myeloid cells. When CD47 binds to SIRPa, it sends inhibitory signals to the macrophage that prevent phagocytosis.

Therapies that block the CD47/SIRPa axis stimulate macrophage engulfment and destruction of cancer cells in preclinical models and show encouraging signs of efficacy in ongoing clinical trials for solid and hematologic malignancies (11-15). However, in many cases, blockade of CD47 by itself is not sufficient to induce macrophage anti-tumor responses, but instead it lowers the threshold for macrophage activation in the presence of a second stimulus (16). Thus, CD47- blocking therapies may work best in combination with other therapies that make cancer cells more vulnerable to macrophage attack (16-18). At present, the best combination modalities remain unknown, and unbiased screening efforts have not been performed to rationally identify drugs that synergize with anti-CD47 agents or other macrophage-directed therapies.

In this study, we aimed to discover novel therapeutic strategies to sensitize lung cancer cells to macrophage-mediated destruction. We developed a high-throughput screening platform to measure anti-tumor function by primary human macrophages in vitro, and we used this system to perform an unbiased screen of 800 FDA-approved drugs. From these efforts, we identified genotype-directed targeted therapies as drugs that make *EGFR* mutant lung cancer cells more vulnerable to macrophage attack. We found these results broadly applied to lung cancers bearing other types of driver mutations and other types of targeted therapies. Our findings illuminate a novel strategy to enhance the efficacy of targeted therapies for lung cancers with oncogenic driver mutations by combining them with anti-CD47 agents.

## RESULTS

### Unbiased drug screens identify EGFR inhibitors as drugs that sensitize cancer cells to macrophage-mediated cytotoxicity

To identify drugs that make lung cancer cells more vulnerable to macrophage-mediated destruction, we developed a novel, unbiased screening platform that measures macrophage anti-tumor function in a high-throughput manner (Figure 1A). We first employed this platform to study *EGFR* mutant lung cancer. We differentiated primary human macrophages ex vivo and co-cultured them in 384-well plates with GFP+ PC9 cells, a human *EGFR* mutant lung cancer cell line (19). A small molecule library of 800 FDA-approved drugs (Supplemental Figure 1) was added to the wells at 5.0 uM concentration along with a CD47-blocking antibody. We then co-cultured the cells for 3-5 days and performed whole-well imaging to quantify the surviving GFP+ area. We evaluated the ability of each drug to kill the GFP+ PC9 cells in the presence of activated macrophages compared to GFP+ PC9 cells alone. From this analysis, we identified two drug classes that specifically inhibit macrophage anti-tumor function (steroids, retinoids), and two drug classes that are less effective when macrophages are present (anthracyclines, other chemotherapy drugs) (Figure 1, B-E, and Supplemental Figure 2 and 3). In contrast, two EGFR tyrosine kinase inhibitors (TKIs)—erlotinib and gefitinib—markedly and specifically enhanced the ability of the macrophages to kill the PC9 cells. These drugs synergized with anti-CD47 therapy and resulted in over 4-fold enhancement of macrophage-mediated cytotoxicity (Figure 1, B-E, and Supplemental Figure 2-4). Given that the PC9 cells contain an activating mutation in *EGFR* (19), we hypothesized that EGFR inhibitors act on the cancer cells to prime them for macrophage mediated-destruction.

**Figure 1:**
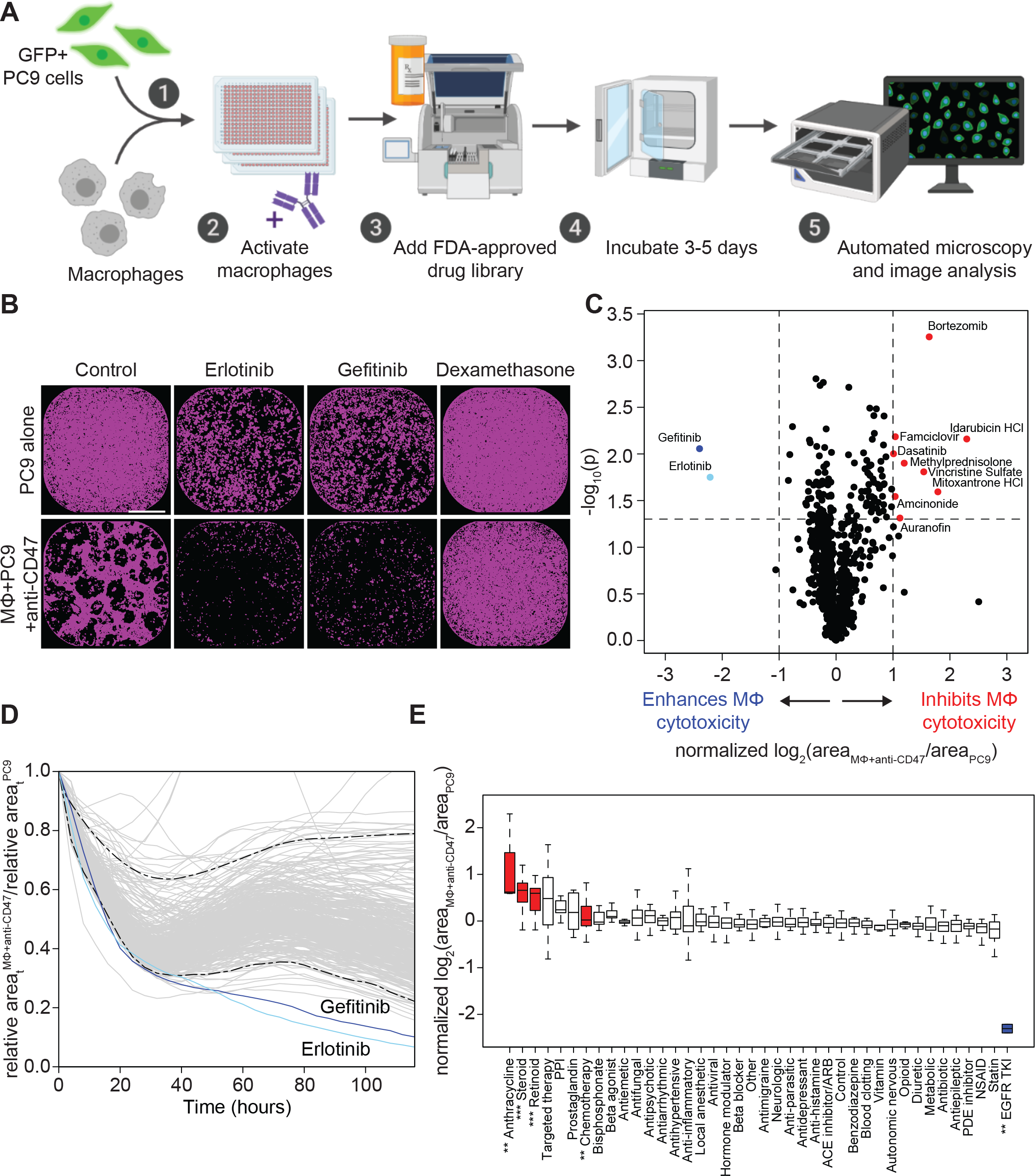
An unbiased compound library screen identifies cooperation between targeted therapy and macrophage-directed immunotherapy for *EGFR* mutant lung cancer. (**A**) Experimental setup of an unbiased functional screen to identify drugs that synergize with macrophage-directed immunotherapy. Primary human macrophages were co-cultured in 384- well plates with GFP+ PC9 cancer cells (a human *EGFR* mutant NSCLC cell line). The wells were treated with 10 ug/mL anti-CD47 antibody, and then a drug compound library (n = 800 FDA-approved drugs) was overlayed at a concentration of 5 uM. The cells were incubated for 3- 5 days and GFP+ area was quantified by automated microscopy and image analysis. As controls, GFP+ PC9 cells were cultured with each individual drug of the library alone for comparison. (**B**) Representative images of whole-well microscopy showing GFP+ area (purple) as quantified by automated image analysis from wells treated with drugs found to enhance (erlotinib, gefitinib) or inhibit (dexamethasone) macrophage-dependent cytotoxicity of PC9 cells. Scale bar, 800 um. (**C**) Volcano plot summarizing the results of the drug library screen. Each point represents the mean of each individual drug treatment condition from n = 5 experimental trials. The phenotypic effect size (x-axis) is depicted as log2 fold-change of GFP+ area in the macrophage+anti-CD47 condition relative to PC9 cells alone. Values were normalized to account for variation due to well position. Dashed lines represent 2-fold change in effect size (x-axis) or p<0.05 by t test (y-axis). Gefitinib and erlotinib (blue) were identified as the top enhancers of macrophage-dependent cytotoxicity of PC9 cells, whereas drugs depicted in red inhibited macrophage dependent cytotoxicity or were drugs that macrophages protected against. (**D**) Representative curves showing macrophage-dependent cytotoxicity over time as represented by decreases in GFP+ area of macrophage+anti-CD47 condition relative to the control condition. Curves from one representative plate showing gefitinib and erlotinib enhance macrophage-dependent cytotoxicity within ∼48 hours. Dashed lines indicate the empirical 95% tolerance interval. (**E**) Box and whisker plot of drug classes included in the screen as ranked by normalized log2 fold-change of GFP+ area in macrophage versus PC9 control condition. Each box indicates the median, interquartile range, maxima and minima (excluding outliers) for the indicated drug class. Drug classes that significantly increased relative GFP+ area are depicted in red, whereas EGFR TKIs (blue) were identified as the only drug class that significantly decreased relative GFP+ area. Each class of drugs was compared with controls (DMSO and empty wells) using a t test (**FDR<0.01, ***FDR<0.001).

### EGFR inhibitors promote macrophage phagocytosis of EGFR mutant lung cancer cells

To investigate the therapeutic potential of combining EGFR inhibitors with anti-CD47 antibodies, we first examined whether CD47 could be a genuine target for lung cancers bearing driver mutations. Using flow cytometry, we evaluated cell-surface expression of CD47 on established and patient-derived cell lines containing *EGFR* or *KRAS* driver mutations or oncogenic *ALK* fusions. We found that CD47 was highly expressed on the cell surface of all specimens tested (Figure 2A). We also compared CD47 expression relative to other surface antigens that regulate macrophage activity, including MHC class I, PD-L1, CD24, and calreticulin (20-23). We found that both CD47 and MHC class I were highly expressed, whereas other macrophage immune checkpoint molecules were low or absent (Figure 2B). We also examined CD47 expression on primary lung cancer cells from malignant pleural effusions and observed high CD47 expression on the cell surface (Figure 2C).

**Figure 2:**
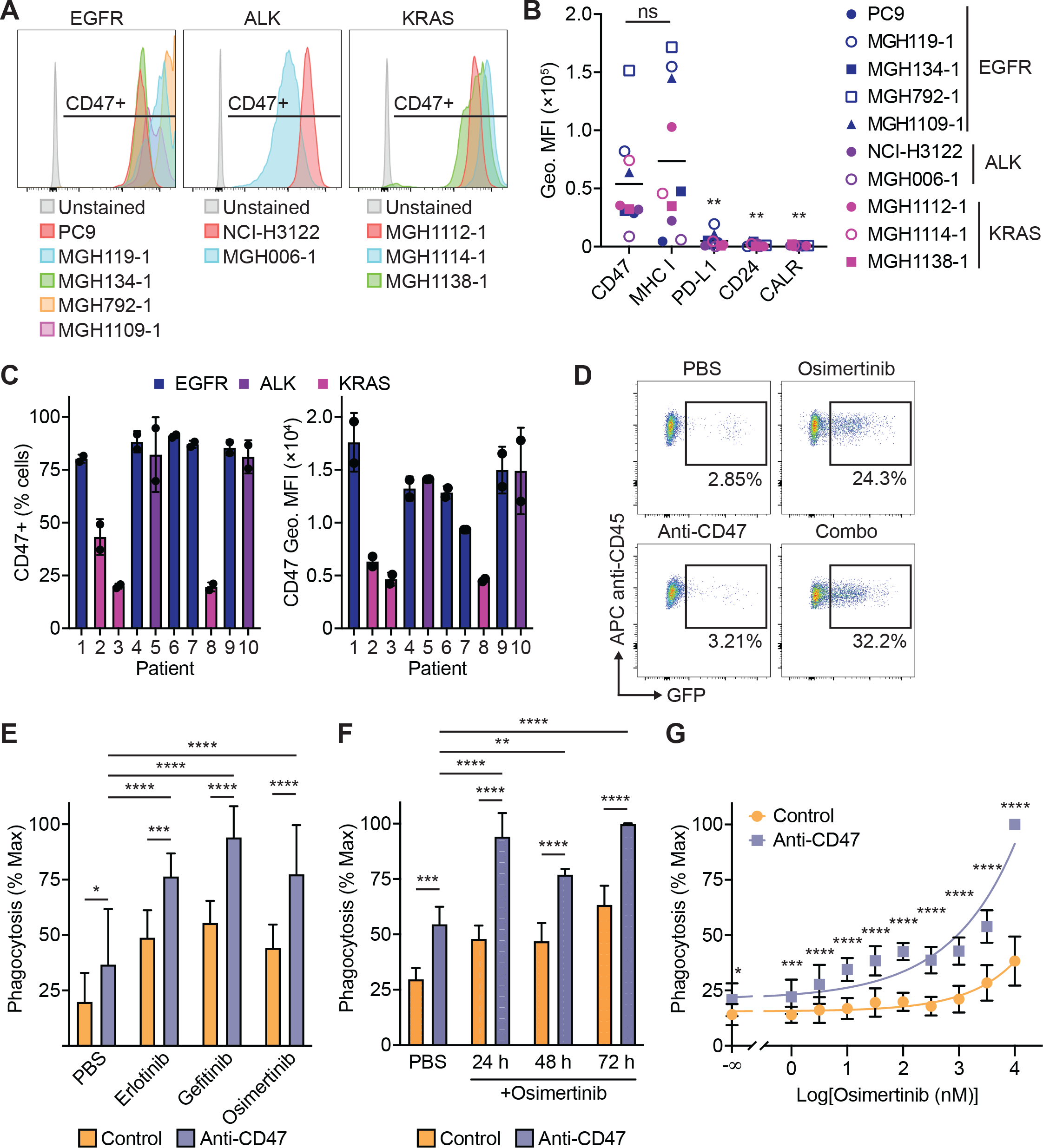
Combined targeting of EGFR and CD47 enhances macrophage phagocytosis in vitro. (**A**) Histograms showing CD47 expression on the surface of NSCLC cell lines containing the indicated driver mutations as assessed by flow cytometry. (**B**) Flow cytometric analysis of macrophage immune checkpoint molecules on the surface of established and patient-derived NSCLC cell lines containing the indicated driver mutations. Geometric mean fluorescence intensity (Geo. MFI) for each antigen was compared to CD47. (**C**) CD47 expression on EpCam+ cancer cells from malignant pleural effusion specimens of patients with NSCLC containing the indicated driver mutations. Left, percent of CD47-positive cells. Right, CD47 geometric MFI. Data represent mean ± SD from 3 technical replicates of n = 10 independent patient specimens. (**D)** Representative examples of phagocytosis assays using primary human macrophages and GFP+ PC9 cells. The PC9 cells were exposed to vehicle control (PBS) or 1 uM EGFR TKI (erlotinib, gefitinib, or osimertinib) for 24 hours. The cells were then collected and co-cultured with primary human macrophages ± an anti-CD47 antibody for 2 hours. Phagocytosis was measured by flow cytometry as the percentage of macrophages containing engulfed GFP+ PC9 cells as indicated in plots. (**E**) Quantification of phagocytosis assays using the indicated EGFR TKIs at 1 uM concentration. Phagocytosis was normalized to the maximal response for each independent donor. Data depict mean ± SD from n = 9 independent blood donors combined from 3 independent experiments using CFSE+ or GFP+ PC9 cells. (**F)** Phagocytosis assays performed using GFP+ PC9 cells exposed to 1 uM osimertinib for varying amounts of time prior to co-culture with primary human macrophages (n = 4 independent donors). The cells were collected and analyzed for phagocytosis as in (F). (**G**) Phagocytosis assays performed using GFP+ PC9 cells exposed to varying concentrations of osimertinib for 24 hours prior to co-culture with human macrophages. Data represent mean ± SD from of 2 independent experiments using a total of n = 8 individual macrophage donors with 3 technical replicates per donor. (B, E-G) ns, *p<0.05, **p<0.01, ***p<0.001, ****p<0.0001 by one-way (B, E) or two-way (F-G) ANOVA with Holm-Sidak multiple comparison test.

We next determined if CD47 could exert a functional role to protect lung cancer cells from macrophage phagocytosis, and whether treatment with EGFR TKIs could enhance phagocytosis. We exposed GFP+ PC9 cells to 1.0 uM TKI (erlotinib, gefitinib, or osimertinib) for 48 hours and then co-cultured the cells with primary human macrophages for 2 hours alone or with a CD47-blocking antibody (24). Regardless of which TKI was used, maximal phagocytosis occurred with the combination of an anti-CD47 antibody and TKI-treated cells (Figure 2, D and E). This effect was maximal within 24 hours of TKI exposure (Figure 2F), a timepoint at which the cancer cells are actively proliferating and only exhibit minimal apoptosis (Supplemental Figure 5). Furthermore, this effect exhibited a dose-response relationship such that greater concentrations of osimertinib resulted in greater macrophage phagocytosis upon anti-CD47 treatment (Figure 2G).

### The combination of EGFR inhibitors and anti-CD47 antibodies eliminates persister cells in vitro

To model the interactions between macrophages and cancer cells as they occur over extended durations of time, we developed ‘long-term’ co-culture assays in which GFP+ cancer cells are co-cultured with primary human macrophages and drug treatments for up to 14 days. We perform whole-well imaging over the co-culture period, and the GFP+ area is quantified over time as a metric of cancer cell growth or death. This experimental system integrates all possible mechanisms of macrophage-mediated cytotoxicity and can evaluate persister cell formation in response to targeted therapies. Using these assays, we co-cultured GFP+ PC9 cells with vehicle control, EGFR TKIs (erlotinib, gefitinib, or osimertinib), anti-CD47, or the combination of anti-CD47 and an EGFR TKI (Figure 3, A-C, Supplemental Figure 6, Supplemental Figure 7, A and B, and Supplemental Movie 1-4). At baseline, macrophages exerted no substantial anti-tumor effect on the PC9 cells. Each individual TKI was able to inhibit the growth of the PC9 cells by themselves, but persister cells always formed and accounted for ∼15% of the cells after 14 days. Treatment with an anti-CD47 antibody caused the formation of patches or foci of cancer cells that remained but also was not able to fully eliminate all cancer cells from the well.

**Figure 3:**
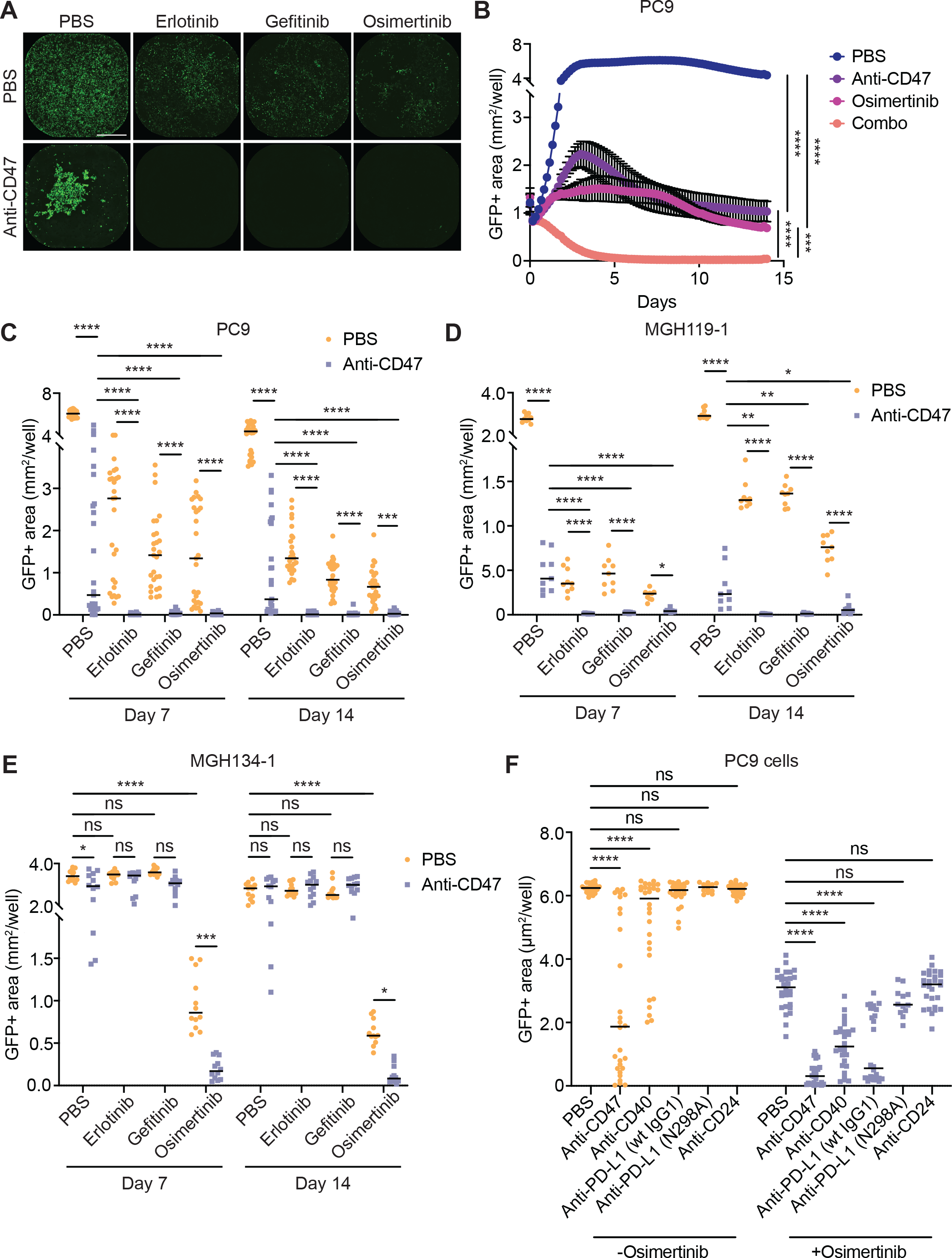
Combining TKIs with anti-CD47 antibodies eliminates *EGFR* mutant persister cells in long-term co-cultures assays with human macrophages. (**A**) Representative images showing quantification of GFP+ fluorescence from co-culture assays on day 6.5. GFP+ PC9 cells were co-cultured in 384-well plates with primary human macrophages alone or with anti-CD47 antibodies (10 ug/mL) and the indicated EGFR TKIs (1 uM). Whole-well microscopy and automated image analysis was performed to quantify GFP+ area (purple) per well over time. Scale bar, 800 um. (B) Representative plot showing growth curves of GFP+ PC9 cells in co-culture with primary human macrophages over time. The cells were treated with 1 uM osimertinib and/or 10 ug/mL anti-CD47 antibody as indicated. Data represent mean ± SEM of three technical replicates per donor using n = 9 independent macrophage donors. Data are combined from two independent experiments with statistical analysis on day 14 of co-culture. (C) Growth of GFP+ PC9 cells in co-culture with primary human macrophages in the absence or presence of anti-CD47 antibodies (10 ug/mL) and the indicated EGFR TKIs (1 uM). Points represent individual replicates, bars represent mean. Data represent three technical replicates per donor using n = 9 independent macrophage donors combined from two independent experiments. (D) Growth of GFP+ MGH119-1 patient-derived cells in co-culture with primary human macrophages in the absence or presence of anti-CD47 antibodies (10 ug/mL) and the indicated EGFR TKIs (1 uM). Points represent individual replicates, bars represent mean. Data represent three technical replicates per donor using n = 3 independent macrophage donors. (E) Growth of GFP+ MGH134-1 patient-derived cells in co-culture with primary human macrophages in the absence or presence of anti-CD47 antibodies (10 ug/mL) and the indicated EGFR TKIs (1 uM). MGH134-1 cells are resistant to first-generation EGFR TKIs (erlotinib, gefitinib) but sensitive to third-generation TKIs (osimertinib). Points represent individual replicates, bars represent mean. Data represent three technical replicates per donor using n = 4 independent macrophage donors. (F) Growth of GFP+ PC9 cells in co-culture assays with primary human macrophages and the indicated macrophage immune checkpoint inhibitors (10 ug/mL). Cells were cultured with antibodies alone or in combination with 100 nM osimertinib. Points represent individual replicates, bars represent means. Data represent GFP+ area on day 6.5 with 3-4 technical replicates per donor and n = 4-8 independent macrophage donors. (B-F) ns, not significant, *p<0.05, **p<0.01, ***p<0. 001, ****p<0.0001 by one-way (B-E) or two-way (F) ANOVA with Holm-Sidak’s multiple comparisons test.

However, the combination of any EGFR TKI with an anti-CD47 antibody dramatically eliminated cancer cells and prevented development of persister cells (Figure 3, A-C, Supplemental Figure 6, Supplemental Figure 7, A and B, and Supplemental Movie 4). These effects were observed over a range of concentrations, with the IC_50_ of the anti-CD47 antibody improving from 223.2 ng/mL (95% CI 158.2-317.3) to 71.25 ng/mL (95% CI 52.39-97.22) upon combination with gefitinib (Supplemental Figure 7B). We observed similar effects using GFP+ MGH119-1 cells, a patient-derived *EGFR* mutant lung cancer cell line (25) (Figure 3D and Supplemental Figure 7C). Again, the combination of each TKI with an anti-CD47 antibody elicited the greatest anti-tumor response and eliminated or prevented the formation of persister cells.

To understand whether the effects of the combination therapy were dependent on sensitivity to a particular TKI, we also tested GFP+ MGH134-1 cells, a patient-derived cell line with a secondary *EGFR^T790M^* mutation (25, 26). This cell line is consequently resistant to erlotinib and gefitinib but sensitive to osimertinib. In co-culture assays, only osimertinib rendered the cancer cells more vulnerable to the anti-CD47 antibody, whereas neither erlotinib or gefitinib inhibited cell growth as single agents nor in concert with CD47 blockade (Figure 3E and Supplemental Figure 7D). These findings suggest that disruption of oncogenic signaling from EGFR is required for enhanced macrophage-mediated cytotoxicity in response to anti-CD47 therapy. Furthermore, they indicate that the TKIs are acting on the cancer cells rather than exerting off-target effects to the macrophages.

We next evaluated whether targeting CD47 was unique compared to other reported macrophage-directed therapies. We tested a panel of antibodies to macrophage immune checkpoints including CD47, CD40, PD-L1, and CD24 (21, 22, 27). We found that as single agents, only the anti-CD47 antibody and an anti-CD40 agonist antibody were able to induce significant macrophage-mediated cytotoxicity of GFP+ PC9 cells (Figure 3F). When combined with osimertinib, the anti-CD47 antibody elicited the greatest anti-tumor response and fully eliminated persister cells (Figure 3F).

### The efficacy of the combination therapy extends to lung cancers with other alterations in the RTK-MAPK pathway

Our results indicate that for *EGFR* mutant lung cancer, disabling signals from EGFR makes the cells more vulnerable to macrophage-mediated destruction. We reasoned that our findings could also apply to lung cancers containing other types of driver mutations. We therefore examined GFP+ NCI-H3122 cells, which contain an oncogenic *EML4-ALK* fusion (28). In co-culture with macrophages, the anti-CD47 therapy significantly impaired the growth of the cancer cells as a single agent (Figure 4A and Supplemental Figure 8). ALK-specific TKIs (crizotinib, alectinib, or lorlatinib) also impaired the growth of the cancer cells as single agents, but a substantial number of persister cells remained in culture. However, the combination of an ALK inhibitor with an anti-CD47 antibody yielded the greatest anti-tumor response, effectively eliminating all cancer cells from the culture (Figure 4A and Supplemental Figure 8). These effects occurred with a dose-response relationship, with the IC_50_ of lorlatinib decreasing from 10.29 nM (95% CI 8.665-12.22) to 2.135 nM (95% CI 0.6934-6.261) with the combination of an anti-CD47 antibody (Figure 4B).

**Figure 4:**
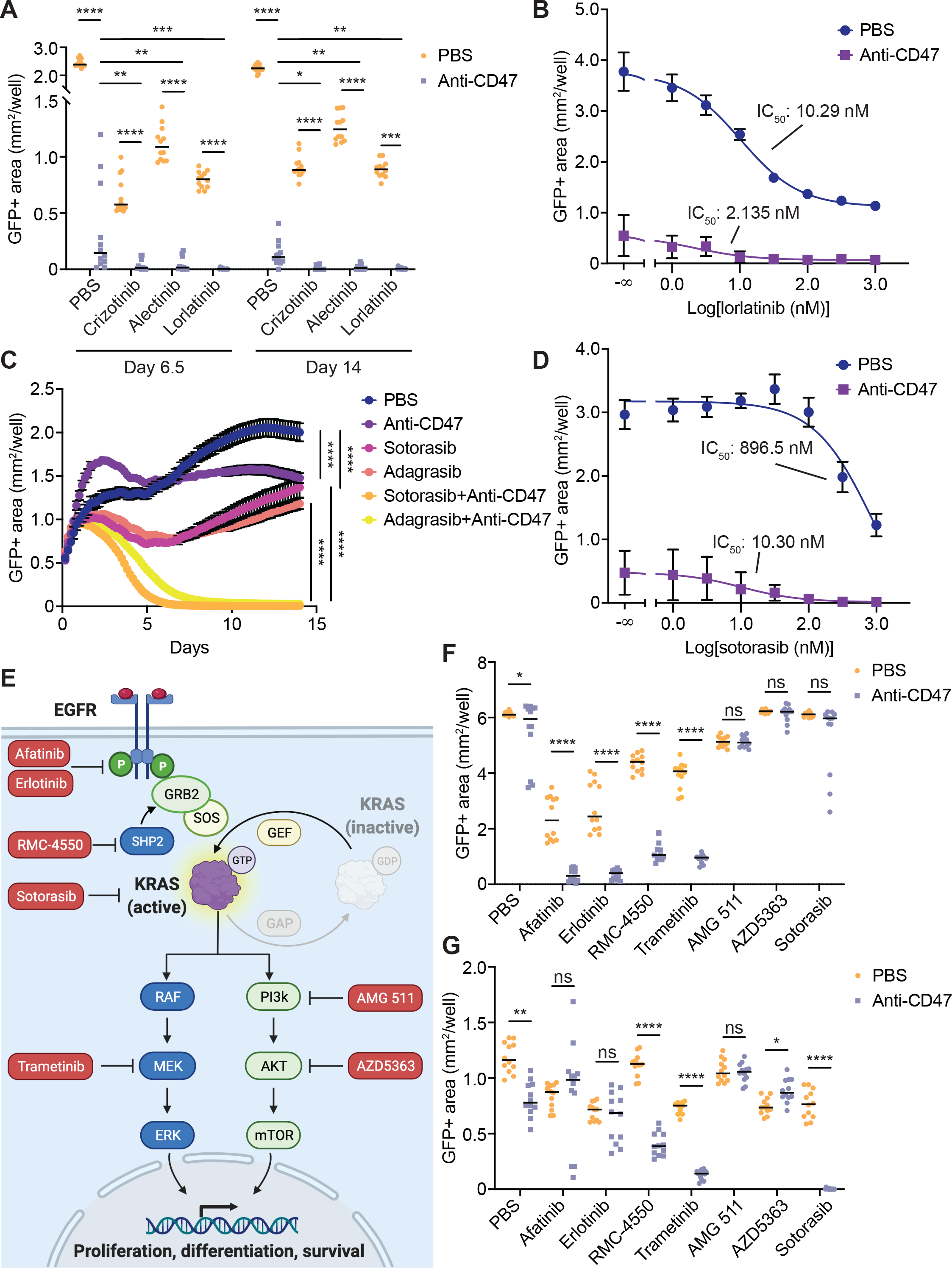
Targeted inhibition of the MAPK pathway primes NSCLC cells for macrophage-mediated destruction. (**A**) Growth of GFP+ NCI-H3122 (a human *ALK*-positive NSCLC cell line) in co-culture with primary human macrophages in the absence or presence of anti-CD47 antibodies (10 ug/mL) and the indicated ALK-specific TKIs (1 uM). Points represent individual replicates, bars represent mean. Data represent three technical replicates per donor using n = 4 independent macrophage donors. (**B**) Growth of GFP+ NCI-H3122 cells in co-culture with primary human macrophages in the absence or presence of anti-CD47 antibodies (10 ug/mL) and varying concentrations of the ALK-specific TKI lorlatinib. Data represent mean ± SD of three replicates each from n = 4 independent macrophage donors on day 6.5 of co-culture. IC_50_ of lorlatinib alone (PBS) = 10.29 nM (95% CI [8.665, 12.22]) versus IC_50_ of lorlatinib+anti-CD47 = 2.135 nM (95% CI [0.6934, 6.261]). (**C**) Growth of GFP+ NCI-H358 (a human *KRAS^G12C^* mutant NSCLC cell line) in co-culture with primary human macrophages in the absence or presence of anti-CD47 antibodies (10 ug/mL) and the indicated KRAS^G12C^-specific inhibitors (1 uM). Data represent mean ± SEM from three technical replicates per donor using n = 4 independent macrophage donors. (**D**) Growth of GFP+ NCI-H358 cells in co-culture with primary human macrophages in the presence or absence of anti-CD47 antibodies (10 ug/mL) and varying concentrations of the KRAS^G12C^-specific inhibitor sotorasib. Data represent mean ± SD of three replicates each from n = 4 independent macrophage donors on day 6.5 of co-culture. IC_50_ of sotorasib alone (PBS) = 896.5 nM (95% CI [558.6, 1697]) versus IC_50_ of sotorasib+anti-CD47 = 10.30 nM (95% CI [2.949, 40.48]). (**E**) Diagram depicting EGFR-RAS-MAPK signaling pathway. KRAS activation leads to bifurcation of signaling via downstream MAPK pathway elements or the PI3K-AKT pathway. Drugs (red boxes) indicate specific inhibitors of pathway elements used in this study. (**F**) Growth of GFP+ PC9 cells in co-culture with primary human macrophages in the absence or presence of anti-CD47 antibodies (10 ug/mL) and varying EGFR-RAS-MAPK or PI3K-AKT pathway inhibitors as indicated in (E). Points represent individual replicates, bars represent mean. Data represent three technical replicates per donor using n = 4 independent macrophage donors on day 6.5 of co-culture. (**G**) Growth of GFP+ NCI-H358 cells in co-culture with primary human macrophages in the presence or absence of anti-CD47 antibodies (10 ug/mL) and varying EGFR-RAS-MAPK or PI3K-AKT pathway inhibitors as indicated in (E). Points represent individual replicates, bars represent mean. Data represent three technical replicates per donor using n = 4 independent macrophage donors on day 6.5 of co-culture. (A, B, F, G) ns, not significant, *p<0.05, **p<0.01, ***p<0. 001, ****p<0.0001 by one-way ANOVA with Holm-Sidak multiple comparisons test.

Similarly, we investigated cell lines with mutations in *KRAS*, one of the most commonly mutated genes in lung cancer (1). We performed co-culture assays using macrophages and NCI-H358 cells, a human lung cancer cell line containing a *KRAS^G12C^* activating mutation (29). In long-term co-culture assays, we found that anti-CD47 antibodies or KRAS^G12C^ inhibitors (sotorasib or adagrasib) inhibited cancer cell growth over time, but generally had only moderate effects as single agents (Figure 4C). In contrast, the combination of the two therapies had a striking effect with dramatic elimination of tumor cells from culture (Figure 4C). As above, we performed titrations of sotorasib and determined that a dose-response relationship existed and that the combination therapy substantially decreased the IC_50_ of sotorasib (896.5 nM, 95% CI 558.6, 1697 nM; versus 10.30 nM, 95% CI 2.949, 40.48 nM) (Figure 4D).

Since EGFR and other receptor tyrosine kinases (RTKs) can transduce growth signals via the MAPK pathway and/or the PI3K/AKT pathway, we tested a panel of inhibitors to dissect the molecular mediators underlying the effects of the combination therapy (Figure 4E). Using co-culture assays with *EGFR* mutant PC9 cells, we found that any active inhibitor of the MAPK pathway could be enhanced by combination with an anti-CD47 antibody, whereas no significant enhancement was observed when combining an anti-CD47 antibody with AKT or PI3K inhibitors (Figure 4F). Similarly, using *KRAS^G12C^* mutant NCI-H358 cells, we found that inhibition of SHP- 2, KRAS^G12C^, or MEK could be enhanced by combining with anti-CD47 therapy, whereas no significant enhancement was observed when combining with PI3K or AKT inhibitors (Figure 4G). These findings suggest that efficacy of the combination therapy is specifically dependent on inhibition of the MAPK pathway rather than alternative signaling pathways.

To understand changes in macrophage activation states as a consequence of the combination therapy, we performed multiparameter flow cytometry, multiplex cytokine analysis, and transcriptional profiling. In co-culture assays, macrophages exhibited increased phagocytosis in response to either targeted therapies or anti-CD47 antibodies as single agents (Supplemental Figure 9). Treatment with the combination therapy elicited upregulation of the M1 markers CD86 and MHC II with relative downregulation of the M2 markers CD163 and CD206 (Supplemental Figure 10). We also found that the combination of targeted therapies and anti-CD47 elicited a unique pro-inflammatory cytokine signature that included significant increases in MIP-1α, RANTES, MCP-4, and TNFα, and a corresponding decrease in VEGF secretion by the cancer cells (Supplemental Figure 11, A and B). Transcriptional profiling identified 50 genes that were significantly upregulated in macrophages upon treatment with the combination therapy.

This gene set included a predominance of genes involved in phagocytosis, cell-cell adhesion, and major drivers of inflammation such as *MYD88* (Supplemental Figure 11C). These profiling experiments emphasize the robust anti-tumor state of the macrophages in response to the combination treatment.

### The combination of targeted therapies and CD47 blockade is effective in mouse tumor models bearing driver mutations

To study the efficacy of the combination therapy in vivo, we first employed xenograft models of human lung cancer. We used NSG mice, which lack functional T-, B- and NK cells but contain macrophages that can be stimulated to attack tumors (16, 24, 30, 31). Importantly, NSG mice have an allele of SIRPa that cross-reacts with human CD47, therefore they have been used as a gold-standard model for evaluating CD47-blocking therapies in vivo (16, 24, 31, 32). We engrafted mice subcutaneously with PC9 cells and allowed tumors to grow to ∼500 mm^3^. Mice were then randomized to treatment with vehicle control, an anti-CD47 antibody, osimertinib, or the combination of osimertinib and the anti-CD47 antibody (Figure 5A). As a single agent, the anti-CD47 antibody produced no significant inhibition of tumor growth.

**Figure 5:**
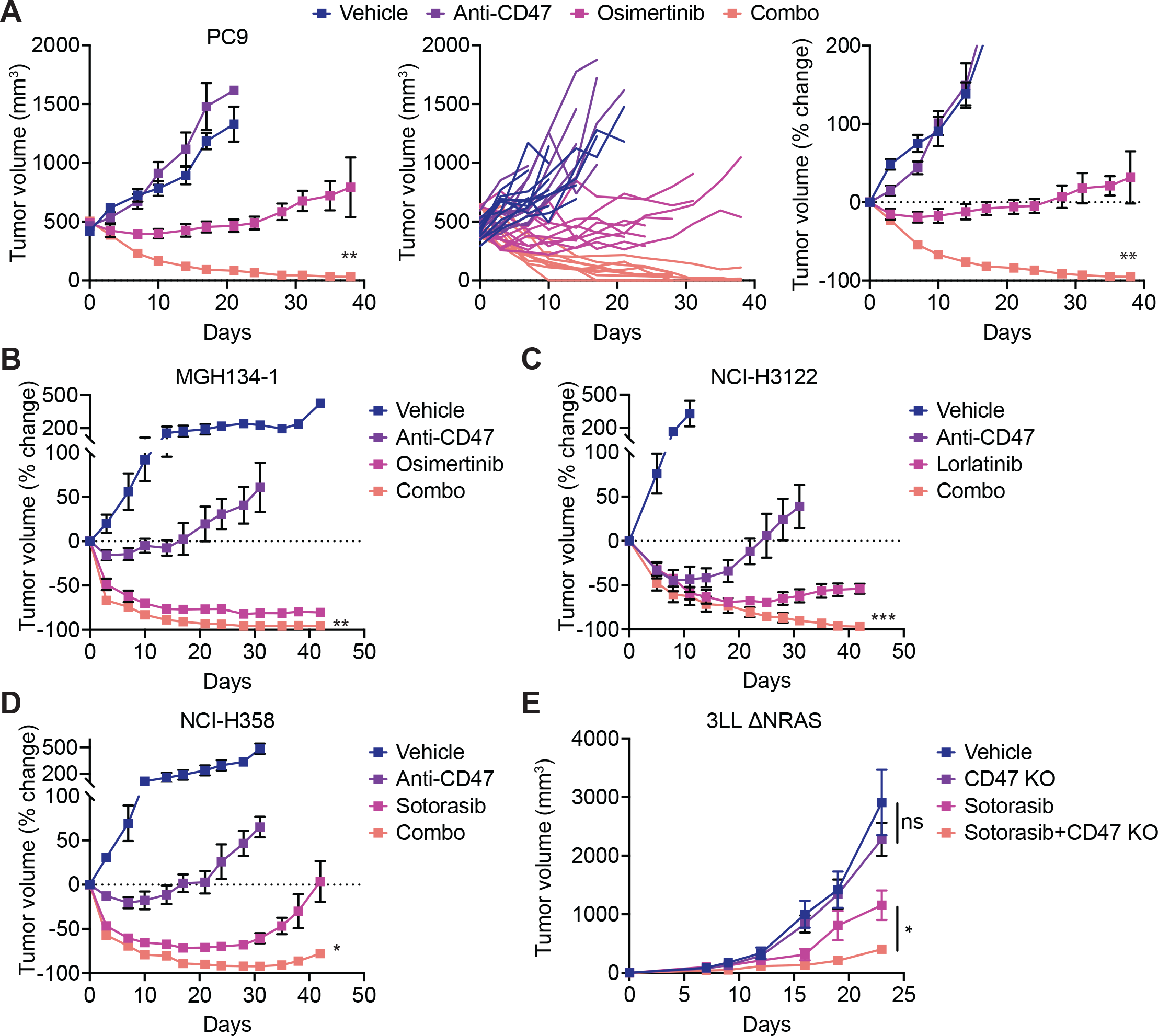
The combination of targeted therapy and CD47 blockade enhances anti-tumor responses in mouse tumor models. (**A**) *EGFR* mutant NSCLC xenograft model of PC9 cells engrafted into NSG mice. Tumors were allowed to grow to approximately 500 mm^3^ and then mice were randomized to treatment with vehicle control, anti-CD47 antibodies (250 ug three times weekly), osimertinib (5 mg/kg five times weekly), or the combination of anti-CD47 plus osimertinib. Tumor volumes were measured over time. Data depict mean tumor volume ± SEM (left), growth curves of individual mice (middle), or percent change in tumor volume from baseline (right). Complete responses were observed in 4/10 mice (40%) in the combination cohort. Data represent n = 9-11 mice per cohort combined from two independent experiments. (**B**) *EGFR* mutant NSCLC xenograft model of MGH134-1 patient-derived cells engrafted into NSG mice and treated as in (A). Data represent percent change in tumor volume from baseline with mean ± SEM of n = 4 mice per cohort. (**C**) *ALK*-positive xenograft model of NCI-H3122 cells engrafted into NSG mice and treated with vehicle control, anti-CD47 antibodies (250 ug three times weekly), lorlatinib (6 mg/kg five times weekly), or the combination of anti-CD47 antibodies and lorlatinib. Data represent percent change in tumor volume from baseline with mean ± SEM of n = 4 mice per cohort. (**D**) *KRAS^G12C^* mutant xenograft model of NCI-H358 cells engrafted into NSG mice and treated with vehicle control, anti-CD47 antibodies (250 ug three times weekly), sotorasib (100 mg/kg five times weekly), or the combination of anti-CD47 antibodies and sotorasib. Data represent percent change in tumor volume from baseline with mean ± SEM of n = 4 mice per cohort. (**E**) Syngeneic tumor model of *KRAS^G12C^* mutant lung cancer using wild-type 3LL ΔNRAS cells or a CD47-knockout variant engrafted into C57BL/6 mice. The mice were treated with vehicle control or sotorasib (30 mg/kg five times weekly) starting on day 7 post-engraftment. Points indicate individual tumor volumes, bars represent median with n = 9-10 mice per treatment cohort. ns, not significant, *p<0.05 by paired t test for the indicated comparisons. (A-D) **p<0.01 by unpaired t test for combo versus targeted therapy.

Treatment with osimertinib as a single agent was able to inhibit tumor growth, but tumors gradually progressed over time. Remarkably, treatment with the combination therapy dramatically reduced tumor burden and elicited complete elimination of tumors in several animals (Figure 5A). We also tested a patient-derived xenograft model of *EGFR* mutant lung cancer (MGH134-1), and again observed the greatest anti-tumor effects from the combination treatment of osimertinib with an anti-CD47 antibody (Figure 5B and Supplemental Figure 12).

To understand whether these findings could extend to other types of lung cancer bearing different driver mutations, we tested models of ALK-positive lung cancer (NCI-H3122 cells) and *KRAS^G12C^* mutant lung cancer (NCI-H358 cells). In each of these models, the greatest anti-tumor effects were observed with the combination of targeted therapy (lorlatinib for NCI-H3122 cells, sotorasib for NCI-H358 cells) and an anti-CD47 antibody (Figure 5, C and D, and Supplemental Figure 12), consistent with our observations in vitro. Similarly, we tested an immunocompetent, syngeneic model of *KRAS^G12C^* mutant lung cancer using 3LL ΔNRAS cells, a variant of Lewis lung carcinoma that harbors an endogenous *KRAS^G12C^* mutation and responds to KRAS inhibitors (33). In this model, we found that genetic ablation of CD47 had no significant effect on tumor growth by itself (Figure 5E and Supplemental Figure 13). Sotorasib was able to inhibit tumor growth as a single agent, yet the greatest inhibition of tumor growth occurred upon sotorasib treatment of a CD47 knockout cell line (Figure 5E). Together, these findings indicate that our in vitro findings translate to in vivo models and that dual blockade of CD47 and oncogenic drivers can enhance anti-tumor responses to lung cancer.

### β2-microglobulin and CD73 are “don’t eat me” signals that can be altered by targeted therapies

To understand the mechanisms by which targeted therapies make cancer cells more vulnerable to macrophage-directed therapies, we generated a panel of seven GFP+ lung cancer cell lines that are each resistant to their respective targeted therapies. The cell lines were generated by prolonged culture in 1 uM of appropriate targeted therapy until resistant cells emerged and grew at rates comparable to their naïve parental counterparts (Figure 6A and Supplemental Figure 14). The lines included PC9 cells (resistant to erlotinib, gefitinib, or osimertinib), NCI-H3122 cells (resistant to crizotinib, alectinib, or lorlatinib), and NCI-H358 cells (resistant to sotorasib). As a consequence of becoming drug resistant, we found that each cell line also became more sensitive to macrophage-mediated cytotoxicity in response to anti-CD47 therapy (Figure 6, B-D). We hypothesized that changes in cell-surface proteins likely mediated this effect, since these proteins are required for intercellular interactions between macrophages and cancer cells. Therefore, we performed comprehensive surface immunophenotyping of naïve parental versus sotorasib-resistant NCI-H358 cell lines to identify differentially expressed surface antigens (Figure 6E). We found two known immunoinhibitory factors, β2-microglobulin (B2M) and CD73 (20, 34), that were substantially downregulated on the sotorasib-resistant line. Moreover, we found that B2M and CD73 were significantly downregulated on additional resistant cell lines, and could be downregulated as a consequence of initial treatment with targeted therapies (Figure 6F and Supplemental Figure 15). To evaluate the functional contributions of these surface proteins, we generated knockout cell lines using CRISPR/Cas9 (Supplemental Figure 12). We found that genetic deletion of B2M could influence macrophage killing in response to anti-CD47 therapy for the majority of cell lines tested (Figure 6G and Supplemental Figure 16). In contrast, CD73 knockout seemed to act as a “don’t eat me” signal only for PC9 cells, and a CD73-blocking antibody was also effective in this setting (Figure 6, H and I, and Supplemental Figure 17). These findings indicate both B2M and CD73 can act as functional macrophage immune checkpoints for lung cancers with driver mutations, and that their downregulation can contribute to vulnerability to macrophage attack. Importantly, individual cancer specimens may differentially rely on these distinct immune checkpoints to evade macrophage-mediated cytotoxicity.

**Figure 6:**
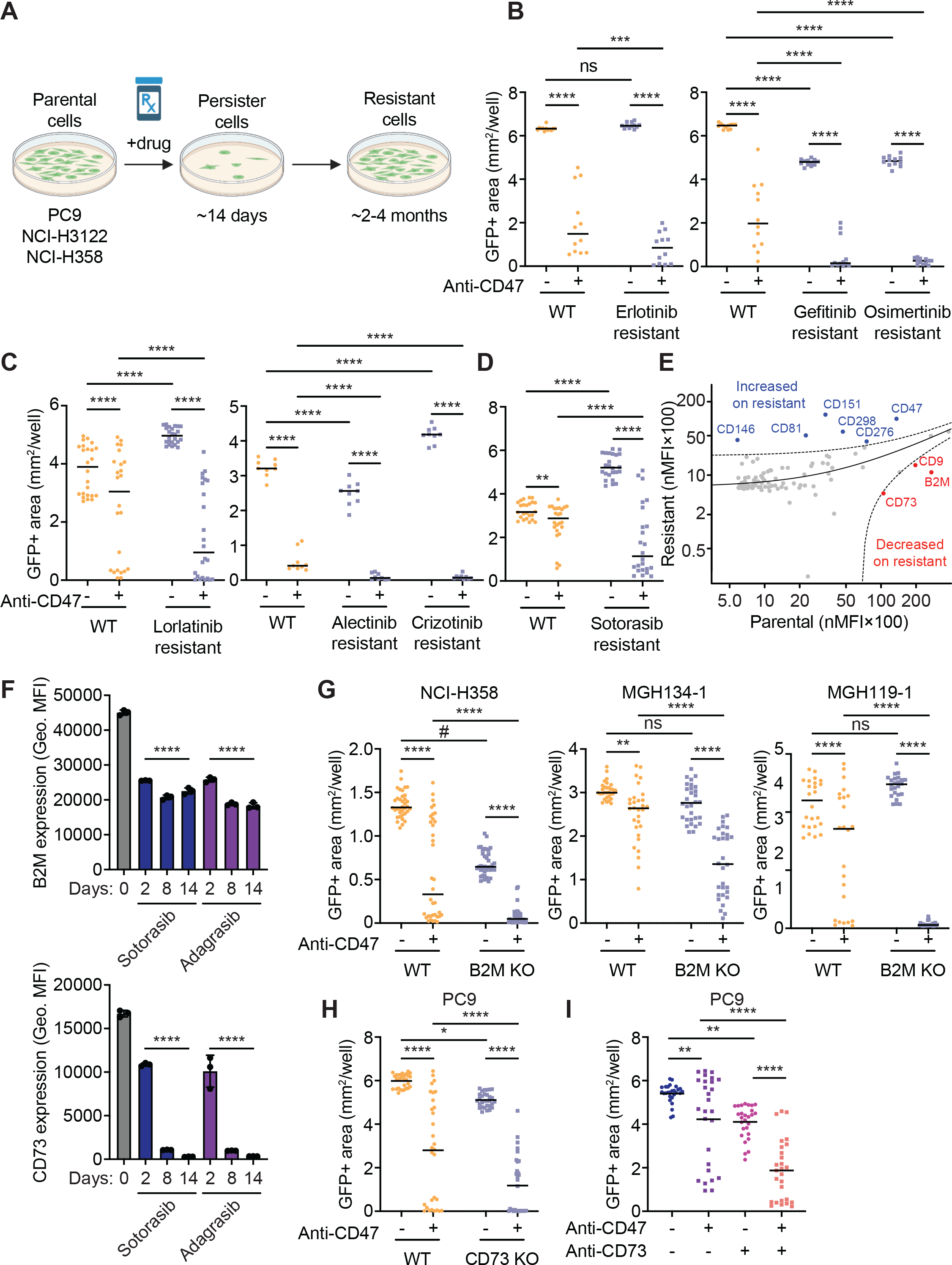
Targeted therapies induce cross-sensitization to anti-CD47 therapy and downregulate B2M and CD73. (**A**) Diagram showing generation of GFP+ cell lines that are resistant to targeted therapies. For each parental cell line (PC9, NCI-H3122, or NCI-H358), cells were cultured in the presence of 1.0 uM of appropriate targeted therapy for prolonged duration until resistant cells emerged and proliferated in culture. (**B-D**) Long-term co-culture assays using GFP+ PC9 cells (B), GFP+ NCI-H3122 cells (C), or GFP+ NCI-H358 cells (D) that are resistant to the indicated targeted therapies. In each case, anti-CD47 therapy resulted in significant enhancement in macrophage-mediated cytotoxicity relative to the naïve parental lines. Experiments performed once with 4 independent donors (erlotinib, gefitinib, osimertinib, alectinib, crizotinib resistant lines) or twice with a total of 8 independent donors (lorlatinib, sotorasib resistant lines) with 3 technical replicates per donor. Bars represent means from analysis at 6.5 days of co-culture. (**E**) Scatter plot showing results of comprehensive surface immunophenotyping of parental NCI-H358 cells versus a GFP+ sotorasib-resistant variant. Each dot represents the normalized mean fluorescence intensity (nMFI) of an individual surface antigen from a total of 354 specificities tested in one experiment. Antigens that exceed the 95% predicted interval for expression on the parental line (red) or resistant line (blue) are indicated. (**F**) Treatment of parental NCI-H358 cells with the indicated targeted therapies causes downregulation of B2M (top) and CD73 (bottom) over time as measured by flow cytometry. ****p<0.0001 for each drug treatment condition compared to time = 0 h by one-way ANOVA with Tukey’s multiple comparison test. (**G**) Evaluation of wild-type versus B2M KO lung cancer cell lines in long-term co-culture assays with human macrophages. (**H**) Evaluation of wild-type versus CD73 KO PC9 cells in long-term co-culture assays with human macrophages. (**I**) Treatment of PC9 cells with a CD73-blocking antibody alone or in combination with anti-CD47 in long-term co-culture assays with human macrophages. (G-I) Data represent at least two independent experiments performed with 6-12 independent macrophage donors. (B-D, G-I) ns, not significant, *p<0.05, **p<0.001, ***p<0.001, ****p<0.0001 by two-way ANOVA with Holm-Sidak multiple comparisons test; #GFP+ area underrepresented due to high confluency of wells and was not visually different by phase microscopy.

## DISCUSSION

To identify therapies that sensitize cancer cells to macrophage-mediated destruction, we performed an unbiased, cell-based functional screen of 800 FDA-approved drugs using primary human macrophages as effectors. We identified genotype-directed targeted therapies as those that prime lung cancer cells for macrophage-mediated destruction. In subsequent in vitro and in vivo validation studies, we found these results extended to multiple NSCLC models harboring diverse oncogenic driver alterations treated with their corresponding targeted therapies. Notably, our findings complement and advance those from a recent study by Hu and colleagues who found KRAS^G12C^ inhibitors could enhance innate immune responses and recruit more macrophages to the tumor microenvironment (35). Interestingly, the effects of the combination therapy may be unique to inhibition of the RTK-MAPK pathway, as similar enhancement was not observed as a class effect of chemotherapy drugs in our screen or when using AKT or PI3K inhibitors. Our findings have immediate translational implications since they suggest the combination of targeted therapies and CD47-blocking therapies could be an optimal strategy for treating patients with lung cancers with driver mutations. Furthermore, we also demonstrate that cell-intrinsic resistance to targeted therapies can cross-sensitize to CD47 blockade.

To date, clinical trials combining targeted therapies and T cell-directed immune checkpoint inhibitors have not been successful for lung cancer. These studies have either been limited in efficacy or demonstrated excessive toxicity. As an example, the combination of osimertinib with an anti-PD-L1 antibody showed severe side effects in nearly 50% of patients (8). Macrophage-directed therapies are an orthogonal treatment modality and may benefit different patients than T cell-directed therapies. This is particularly true since macrophages have inherent ability to kill cancer cells when provided with an appropriate stimulus, whereas T cell cytotoxicity is intertwined with the tumor mutational burden and the presence of neoantigens. Interestingly, we found that downregulation of B2M and CD73 could contribute to enhanced sensitivity to macrophage killing. B2M is required for MHC class I expression on the cell surface, which CD8 T cells depend on for antigen recognition. However, B2M also acts as a “don’t eat me” signal by binding to LILRB1, an inhibitory receptor on macrophages (20). Downregulation of B2M could decrease antigen presentation to make cancer cells resistant to T cell-directed immunotherapies while simultaneously making them more vulnerable to macrophage attack.

Thus, B2M expression may reflect a critical pivot point between innate and adaptive immune activation. Similarly, CD73 is an ectoenzyme that catalyzes the breakdown of AMP to immunosuppressive adenosine in the tumor microenvironment (34). Its downregulation may make lung cancer cells more sensitive to macrophage-mediated cytotoxicity. Our findings suggest lung cancer specimens may differentially rely on these immunosuppressive signals to evade detection by macrophages.

The high-throughput screening platform we developed is a robust system to identify drugs that activate or inhibit macrophage-mediated cytotoxicity. Of note, our screen also identified putative drug-drug interactions that may negatively affect CD47-blocking therapies or other macrophage-directed therapies for cancer. For example, steroids and retinoids were classes of drugs that were generally inhibitory to macrophages and abrogated the effects of anti-CD47 therapy. Moreover, anthracyclines were effective at killing cancer cells by themselves as single agents, but they were completely inactive in the presence of macrophages. These findings warrant further investigation in vivo, but they suggest these drug classes should be limited or avoided when treating patients with macrophage-directed therapies. In the future, our screening platform could be applied to high-throughput investigation of larger compound libraries. Since the platform uses functional interrogation of primary human macrophages, it is target agnostic and therefore offers great opportunity to understand fundamental biology and discover new drug candidates.

As limitations of our study, our experimental approach focused on in vitro studies using primary human macrophages, xenograft mouse models, and syngeneic immunocompetent mouse models. Although the combination of these studies has been useful to predict the safety and efficacy of CD47-blocking therapies in clinical trials, all preclinical models have inherent limitations and it is possible that our findings do not translate to patients as expected. As another limitation, toxicity has been observed with some anti-CD47 agents, primarily relating to on-target toxicity to normal, healthy red blood cells that express CD47 on their surface (15, 36, 37). Targeted therapies for lung cancer generally do not have significant effects on red blood cells (2, 3, 5), therefore we do not expect enhanced toxicity from the combination strategy.

However, this possibility should be evaluated and monitored further in clinical trials. Finally, our study focused exclusively on lung cancers with driver mutations. We expect our findings will extend to other types of cancers, particularly those with RTK-MAPK pathway mutations, but further experimentation is needed to investigate these principles.

In order to translate our discoveries to patients, investigation of the combination strategy in clinical trials is warranted. There exist multiple FDA-approved targeted therapies for lung cancer (4), and multiple CD47-blocking therapies are now under investigation in clinical trials (38). Thus, combining these two treatment strategies is feasible and can occur in the near term. Our data suggest the most effective setting to test the combination strategy would be in patients with lung cancers with oncogenic driver mutations that are naïve to targeted therapy. However, our data also suggest CD47-blocking therapies may be effective as single agents in patients who have progressed on targeted therapy. Overall, our findings indicate that combining targeted therapies with CD47-blocking agents or other macrophage-directed therapies may be an ideal way to merge the fields of precision medicine and immuno-oncology for the benefit of patients.

## METHODS

### Cell lines

PC9, NCI-H3122, NCI-H358 cells were obtained from the Hata Laboratory (Massachusetts General Hospital). Cell line identities were confirmed by STR profiling. Mycoplasma testing was performed prior to banking frozen stocks used in this study and when any uncertainty in culture conditions arose. Patient-derived cell lines (MGH119-1 (25), MGH134- 1 (26), MGH792-1 - *EGFR* mutant NSCLC; MGH006-1 – *ALK*-positive NSCLC; MGH1112-1, MGH1114-1, MGH1138-1 – *KRAS^G12C^* NSCLC) were generated using previously described methods (26). Prior to cell line generation, the patients signed informed consent to participate in a Dana Farber/Harvard Cancer Center Institutional Review Board-approved protocol giving permission for research to be performed on their sample. Human NSCLC cell lines were cultured in in RPMI (Thermo Fisher) supplemented with 10% ultra-low IgG fetal bovine serum (Thermo Fisher), 100 units/mL penicillin, 100 ug/mL streptomycin, and 292 ug/mL L-glutamine (Thermo Fisher). 3LL ΔNRAS cells were provided by the lab of Dr. Julian Downward (The Francis Crick Institute, London, UK) and were cultured in DMEM (Thermo Fisher) supplemented with 10% ultra-low IgG fetal bovine serum, 100 units/mL penicillin, 100 ug/mL streptomycin, and 292 ug/mL L-glutamine. Cell lines were maintained in humidified incubators at 37 degrees C with 5 % carbon dioxide.

GFP+ lines were generated by lentiviral transduction of cell lines using CMV-GFP-T2A- Luciferase pre-packaged virus (Systems Bio). Transduced cells were then sorted for stable GFP expression.

### Genetic modification of cell lines

Knockout cell lines were generated by CRISPR/Cas9-mediated genome editing. A CD47 knockout variant of 3LL ΔNRAS cells was generated using Gene Knockout Kit version 2 targeting murine CD47 (Synthego). Knockout variants of PC9, NCI-H358, MGH119, and/or MGH134 were generated using Gene Knockout Kit version 2 targeting human B2M or human CD73 (Synthego). Gene knockout was performed via ribonucleoprotein transfection with recombinant Cas9 (Synthego). Cells were then stained for surface antigen expression and sorted using a FACSAria II (BD Biosciences) to generate negative cell lines. For murine CD47, staining was performed using APC anti-murine CD47 clone miap301 (BioLegend) and sorting was used to generate a clonal population. For human lines, staining wass performed with APC anti-human B2M clone 2M2 (Biolegend) or APC anti-human CD73 clone AD2 (Biolegend) and used to sort polyclonal lines that were negative for surface antigen expression.

### Ex vivo generation of primary human macrophages

Primary human macrophages were generated from peripheral blood mononuclear cells as previously described (16). Briefly, leukocyte reduction chambers were obtained from healthy human donors from discarded apheresis products via the Crimson Core Biobank (Boston, MA). Monocytes were labeled using StraightFrom Whole Blood CD14 MicroBeads (Miltenyi Biotec) and purified using an autoMACS Pro Separator (Miltenyi Biotec). Monocyte were then cultured in IMDM (Thermo Fisher) supplemented with 10% ultra-low IgG fetal bovine serum, 100 units/mL penicillin, 100 ug/mL streptomycin, and 292 ug/mL L-glutamine and 20 ng/mL human M-CSF (Peprotech) for at least 7 days. Cells were passaged or replated as necessary and typically maintained in culture for 2-4 weeks.

### FDA drug library screen

GFP+ PC9 cells and primary human macrophages were co-cultured in 384-well plates in IMDM supplemented with 10% ultra-low IgG fetal bovine serum, 100 units/mL penicillin, 100 ug/mL streptomycin, and 292 ug/mL L-glutamine and 20 ng/mL human M-CSF. Purified anti-CD47 antibody clone B6H12 (BioXCell or eBioscience) was added at a working concentration of 10 ug/mL. Duplicate control plates were plated with GFP+ PC9 cells alone. A curated library of 800 FDA-approved drugs was transferred to the plates via Echo Acoustic Liquid Handler (Koch Institute High-Throughput Sciences Facility). Cells were then incubated for 3-5 days. Wells were imaged using automated fluorescence microscopy with an Incucyte S3 system (Sartorius). Automated image analysis was performed using Incucyte Analysis Software (Sartorius) to quantify the percentage of GFP+ area occupied by target cells in each well. To assess macrophage-dependent cytotoxicity, GFP+ area in experimental wells (containing GFP+ PC9 cells, macrophages, and anti-CD47 antibodies) was normalized to GFP+ area in control wells (containing GFP+ PC9 cells alone). Library screens were independently repeated n = 5 times, each time using macrophages derived from individual donors. For individual compounds, p-values were computed using a one-sample t test after controlling for row, column and plate effects (39).

### FACS analysis

Analysis of cell surface antigens was performed using live cells in suspension. Cells were blocked with 100 ug/mL mouse IgG (Lampire Biological Laboratories) or Human TruStain FcX (BioLegend). The following antibodies were used for FACS analysis: APC-conjugated anti-CD47 clone B6H12 (Thermo Fisher), Alexa Fluor 647-conjugated anti-HLA-A,B,C (MHC class I) clone W6/32 (BioLegend), APC-conjugated anti-PD-L1 clone 29E.2A3 (BioLegend), Alexa Fluor 647-conjugated anti-CD24 clone ML5 (BioLegend), PE-conjugated anti-CALR clone FMC 75 (Abcam), APC anti-CD45 clone 2D1 or clone HI30 (BioLegend), APC anti-B2M clone 2M2 (BioLegend), APC anti-CD73 clone 2A2 (BioLegend). Viability of cell lines was assessed by staining with 100 ng/mL DAPI (Milipore Sigma). For analysis of primary pleural fluid specimens, Alexa Fluor 488-conjugated anti-EpCam clone 9C4 (BioLegend) was used to mark the malignant cell population. Cells were analyzed using an LSR Fortessa equipped with a High Throughput Sampler (BD Biosciences). Comprehensive surface immunophenotyping was performed using LEGENDScreen Human PE Kit containing 371 pre-titrated antibodies and 354 unique specificities (BioLegend). The experimental run was multiplexed using GFP+ NCI-H358 cells resistant to sotorasib combined with non-fluorescent, parental NCI-H358 cells.

Compensation was performed prior to analysis on an LSR Fortessa using FACSDiva software (BD Biosciences).

### Targeted therapies

The following targeted therapies were used in this study: erlotinib, gefitinib, osimertinib, crizotinib, alectinib, lorlatinib, sotorasib, adagrasib (Selleckchem). MAPK pathway inhibitors included: afatinib, erlotinib, RMC-4550, sotorasib, trametinib, AMG 511, AZD5363 (Selleckchem).

### Phagocytosis assays

In vitro phagocytosis assays were performed as previously described (16). Briefly, GFP+ PC9 cells or CFSE+ PC9 cells were used as target cells. Labeling with CFSE (Thermo Fisher) was performed according to manufacturer’s instructions. Target cells were exposed to EGFR TKIs (erlotinib, gefitinib, or osimertinib) for 24-72 hours. Live, adherent cells were then collected, washed, and co-cultured with primary human macrophages at a target:macrophage ratio of 4:1. Cells were co-cultured in the presence or absence of 10 ug/mL purified anti-CD47 antibody clone B6H12. Cells were co-cultured for 2 hours in serum-free IMDM in round-bottom ultra-low attachment 96-well plates (Corning). After the incubation period, cells were washed and analyzed by flow cytometry. Macrophages were identified using APC anti-CD45 clone 2D1 or clone HI30 and target cells were identified by CFSE or GFP fluorescence. Phagocytosis was quantified as the percentage of macrophages that contained CFSE or GFP+ signal. Phagocytosis was normalized to the maximal response by each independent macrophage donor. Dose-response curves were generated using Prism version 9.2.0 (GraphPad).

### Long-term co-culture assays

Long-term co-culture of primary human macrophages was performed using GFP+ target cells. Cells were cultured in phenol-red free IMDM (Thermo Fisher) supplemented with 10% ultra-low IgG fetal bovine serum, 100 units/mL penicillin, 100 ug/mL streptomycin, and 292 ug/mL L-glutamine and 20 ng/mL human M-CSF. Purified anti-CD47 antibody clone B6H12 was added at a working concentration of 10 ug/mL. Targeted therapies were added at a final working concentration of 1.0 uM or as indicated otherwise. Cells were then co-cultured for 7-14 days with whole-well imaging of phase contrast and GFP channels performed every 4 hours using an Incucyte S3 system. Automated imaging analysis was performed using Incucyte Analysis Software. Images taken on day 6.5 was used as a standard reference time point for data comparison and statistical analysis. For dose-response experiments, sigmoidal dose-response curves were generated using Prism version 9.2.0 (GraphPad) to determine IC_50_ parameters.

### Macrophage immune checkpoint experiments

In long-term assays to assess macrophage immune checkpoint targeting, the following macrophage-directed therapeutics were used: purified anti-CD47 clone B6H12, anti-CD40 clone G28.5 (BioXCell), anti-hPD-L1-hIgG1 (Invivogen), anti-hPD-L1-hIgG1 (N298A) (Invivogen), purified anti-CD24 clone SN3 (GeneTex), and purified anti-CD73 clone AD2 (BioLegend).

### In vivo treatment models

NOD.Cg-Prkdc^scid^ Il2rg^tm1Wjl^/SzJ (NSG) mice (Jackson Laboratory) were used for all xenograft tumor models. Cells were implanted subcutaneously into the flank of NSG mice (age 6-8 weeks) in 0.2 ml 50% BD Matrigel Basement Membrane Matrix in PBS. Tumor volumes were measured twice weekly using electronic calipers and calculated using the formula: mm^3^ = 0.52 × L × W^2^. Mice were randomized to treatment cohorts when tumor volumes reached approximately 500 mm^3^. Drugs used for treatment included: osimertinib (5 mg/kg via oral administration five times per week), lorlatinib (6 mg/kg via oral administration five times per week), sotorasib (100 mg/kg via oral administration five times per week), or anti-CD47 antibody clone B6H12 (250 ug via intraperitoneal injection three times per week). The mice were euthanized when the tumor size exceeded 20 mm in any one dimension or when mice reached humane experimental endpoints according to the Institutional Animal Care and Use Committee-approved protocols.

For a syngeneic, immunocompetent tumor model, C57BL/6 mice (Jackson Laboratory) were engrafted with 3LL ΔNRAS cells or a CD47-knockout variant. Mice were engrafted in the subcutaneous tissue using a dual-flank model. Wild-type tumors were engrafted on one flank, and CD47-knockout tumors were engrafted on the contralateral flank. Mice were then randomized to treatment with vehicle control or sotorasib (30 mg/kg via oral administration five times per week). Tumor volumes were measured twice weekly using electronic calipers and calculated using the formula as above.

### Statistics

Data analysis of FDA drug screens was performed across 5 replicates of the experiments as follows. GFP+ area was quantified from all wells as described above. After logarithmic transformation, row, column and plate effects were controlled using Tukey’s median polish, after which p-values for individual compounds (Fig. 1C) were computed using a one-sample t test (39). For plate-based time-series measurements (Fig. 1D), “relative area” was defined as the GFP+ area at time t divided by the GFP+ area at time zero, obviating the need for controlling row, column and plate effects. To assess the effect of combining macrophage+anti-CD47 conditions with different classes of drugs (Fig. 1E), each class was compared with control wells (i.e. empty or DMSO) using a two-sample t test with the Benjamini-Hochberg method (40) to control for multiple hypothesis testing.

In general, flow cytometry data was analyzed by comparing geometric mean fluorescence intensity (geometric MFI) of individual surface antigens by one-way ANOVA with Holm-Sidak multiple comparison test. Comprehensive surface immunophenotyping was analyzed from a single set of measurements. Data for naïve parental versus sotorasib-resistant lines were uniformly adjusted by linear transformation to correct for negative values. Data were fit using linear regression which was used to define the 95% prediction interval.

In vitro phagocytosis assays were analyzed by comparing the mean percentage of GFP+ macrophages between different conditions. Experiments were performed at least 2 independent trials using a total of 6-8 individual donors except as otherwise indicated. The mean values for relevant comparisons were assessed by two-way ANOVA with Holm-Sidak multiple comparison test using Prism version 9.2.0 (Graphpad).

Long-term co-culture assays were performed by measuring GFP+ area. Data was analyzed by comparing mean GFP+ area between relevant conditions using one-way ANOVA with Holm-Sidak multiple comparison test. Experiments were performed in two independent trials using a minimum of 6-8 donors except as otherwise indicated. IC_50_ values were calculated by fitting data to sigmoidal dose-response curve using Prism version 9.2.0 (Graphpad).

Analysis of mouse xenograft tumor models was performed by unpaired t test of targeted therapy cohorts versus combination therapy cohorts on the last day of tumor measurements using Prism version 9.2.0 (Graphpad). For dual-flank model of 3LL ΔNRAS, statistical analysis was performed by comparing median tumor volume by paired t test using Prism version 9.2.0 (Graphpad).

For all experiments, p<0.05 was considered statistically significant.

### Study approval

The use of primary lung cancer specimens, patient-derived cell lines, and patient-derived xenografts was performed with specimens collected from patients with written informed consent prior to participation under protocols approved by the Dana-Farber/Harvard Cancer Center Institutional Review Board. Experiments with primary human macrophages were exempt from review by the MIT Committee on the Use of Humans as Experimental Subjects since the cells derived from discarded blood products from anonymous donors. All animal studies were conducted in accordance with the guidelines as published in the Guide for the Care and Use of Laboratory Animals and were approved by the Institutional Animal Care and Use Committee of Massachusetts General Hospital or the Massachusetts Institute of Technology Committee on Animal Care.

## Author contributions

Conceptualization: KW, ANH

Methodology: KV, JA, AM, SR, JV, TW, DY, GWB, NP, ANH, KW

Investigation: KV, JA, SR, NP, TW, KW

Visualization: KV, JA, AM, SR, JV, TW, GWB, ANH, KW

Funding acquisition: GWB, KW, ANH

Project administration: KV, JA, GWB, KW, ANH

Supervision: GWB, KW, ANH

Writing – original draft: KW

Writing – review & editing: KV, JA, AM, SR, JV, TW, DY, GWB, ANH, KW

The order of co-first authors was designated based on the duration of time each author contributed to this study.

## Acknowledgments

We thank members of the Weiskopf, Hata, and Weinberg labs for experimental assistance, reagents, and insightful discussions. We thank Donna Hicks, Ferenc Reinhardt, Elinor Eaton, Joana Liu Donaher, Nicholas Polizzi, Beverly Dobson for support. We thank Dr. Robert Weinberg for experimental guidance and insight, and critical review of the manuscript. We thank Dr. Irving Weissman and Dr. Julian Downward for providing cell lines and reagents. We thank Patrick Autissier and the Whitehead Institute Flow Cytometry Core Facility; Jennifer Love, Stephen Mraz, Thomas Vokert and the Genome Technology Core Facility; and the Bioinformatics and Research Computing Core Facility. We thank Jaime Cheah, Christian Soule, and the Koch Institute High Throughput Sciences Facility. Experimental diagrams were created using Biorender.

This work was supported by the Valhalla Foundation (KW), NIH T32 CA09172 (KW, AM), A Breath of Hope Lung Foundation Peg Fisher-Jullie Fight for Life Award (KW), ASCO Conquer Cancer Foundation Young Investigator Award (KW), AACR-AstraZeneca Career Development Award for Physician-Scientists in Honor of José Baselga (KW), Fast Grants, Emergent Ventures, George Mason University (KW), Society for Immunotherapy of Cancer, Holbrook Kohrt Cancer Immunotherapy Translational Memorial Fellowship (KW), American Lung Association Catalyst Award (KW), Anonymous Grant (KW), Dr. Richard Reisman in honor of Jane Reisman (KW), Damon Runyon Cancer Research Foundation Postdoctoral Fellowship (DY), NIH R01 CA137008 (ANH), Ludwig Center at Harvard (ANH), and the Lungstrong foundation (ANH).

## SUPPLEMENTAL MATERIALS

### List of Supplemental Materials

Supplemental Methods

Supplemental Figures 1-17

Supplemental Movies 1-4

### Supplemental Methods

#### Cytokine analysis

Primary human macrophages were co-cultured with PC9 or NCI-H358 cells for 4-7 days. Supernatants were collected and frozen at -80C. Supernatants were subjected to addressable laser bead immunoassay analysis (ALBIA) using a Human Cytokine 71-Plex Discovery Assay (Eve Technologies). Cytokine assay levels were floored (for ″OOR <″ measurements) to 0, and high values (″OOR >″) were replaced with the highest observed concentration. Cytokines with a mean level < 1 were excluded. Statistical analysis to identify cytokines that differed between all targeted therapy+anti-CD47 combination treatment specimens versus all single-agent anti-CD47 specimens was performed with 2-way ANOVA for each cytokine, and p-values were corrected for FDR across all cytokines. Replicate assays were median-summarized. For each combo, the log2(combo / vehicle) ratio was calculated, and the cytokines were sorted by decreasing mean ratio for heatmap representation.

#### Myeloid transcriptional profiling

Primary human macrophages and NCI-H358 cells were co-cultured for 4 days, then cell specimens were collected in QIAzol and RNA was extracted by RNeasy Kits (Qiagen) according to the manufacturer’s instructions. RNA samples were processed using an nCounter SPRINT Profiler from Nanostring Technologies, according to manufacturer’s directions. Briefly, 50 ng of total RNA derived from co-cultured cells was hybridized with the nCounter Myeloid Innate Immunity Panel at 65°C for 18 hours, before being processed on a SPRINT cartridge. The gene counts were subsequently analyzed using Nanostring’s nSolver software. Gene counts were normalized with DESeq2. To identify genes that were consistently different between the combo treatment and other conditions, we used one-way ANOVA, followed by Dunnett’s test. The p-values (the maximum for each gene) were subsequently corrected for FDR. All genes with FDR < 0.05 were mean-summarized across each condition, and the resulting matrix was log2-transformed, mean-centered, and clustered by gene (with uncentered correlation and pairwise average-linkage) for heatmap representation.

---

**Supplemental Figure 1:**
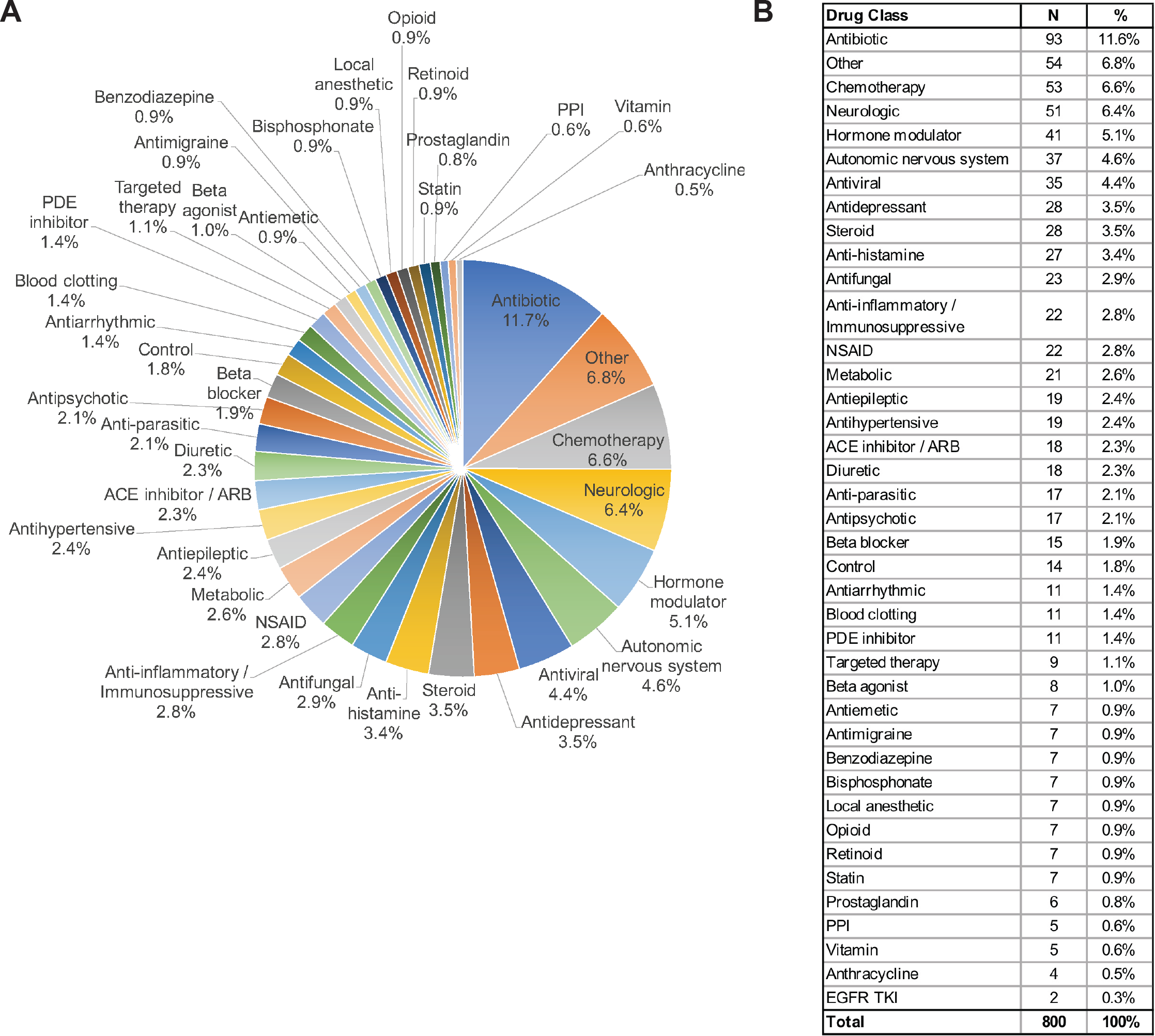
The composition of an FDA-approved drug library used for screening efforts. (**A**) Chart depicting drug classes included in the screening library. Percentages indicate number of drugs per class from a total of *N* = 800 individual drug wells and including 14 DMSO controls. (**B**) Table depicting the number and percentage of drugs from each class included in the screening library.

**Supplemental Figure 2:**
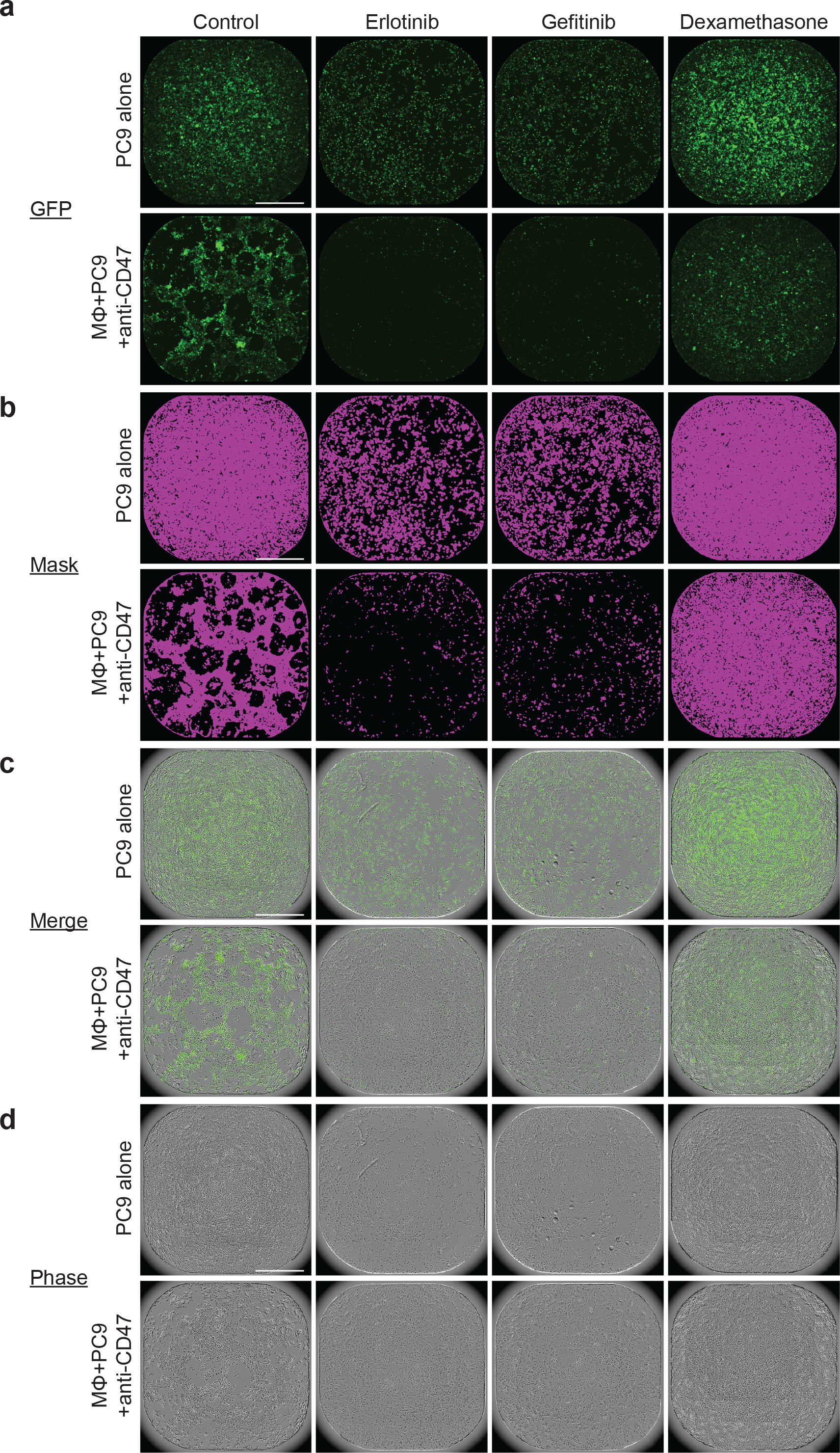
Representative images of wells from small molecule screen using FDA-approved drug library. GFP+ PC9 cells were combined with primary human macrophages and the indicated drug therapies in 384-well plates. Representative images are shown from a single experimental run of the full FDA-approved drug library using macrophages derived from an individual blood donor at t = 3d 16h. (**A**) Whole well imaging of the GFP+ channel from wells treated with the indicated therapies. Erlotinib and gefitinib were identified as drugs that enhance macrophage-dependent cytotoxicity of PC9 cells, while dexamethasone and other steroid compounds were identified as inhibitors of macrophage-dependent cytotoxicity. (**B**) Image mask of GFP+ pixels (purple) used for quantification and analysis. (**C**) Overlay of GFP+ channel with phase contrast imaging. (**D**) Phase contrast imaging showing confluency of wells with GFP+ PC9 cells and primary human macrophages present.

**Supplemental Figure 3:**
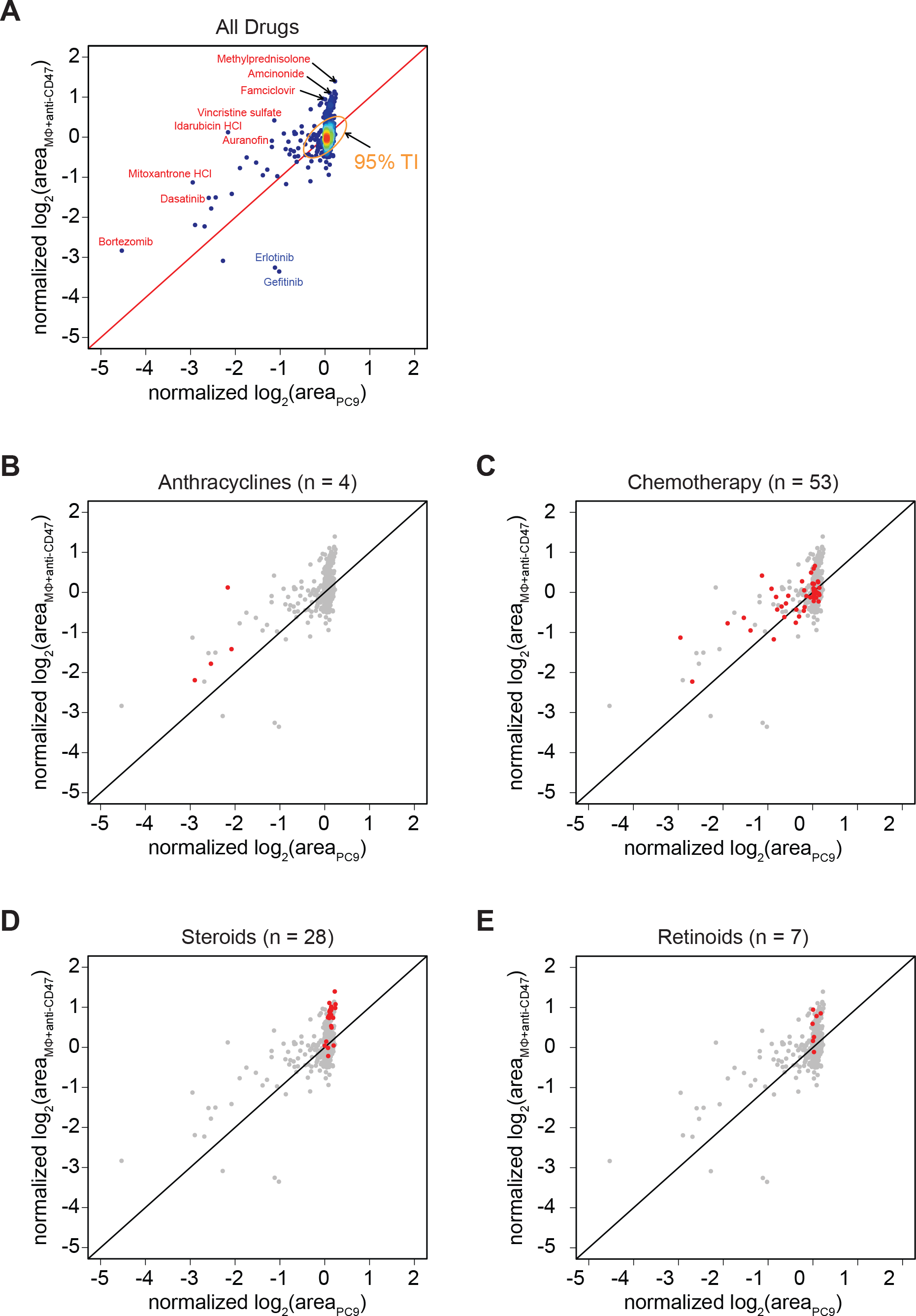
Analysis of high-throughput screen reveals differential activity of drugs from the FDA-approved library. (**A**) Scatter plot showing how drugs affect growth of GFP+ PC9 cells alone (x-axis) versus when they are co-cultured with macrophages and anti-CD47 therapy (y-axis). Points are colored by density from low (blue) to high (red), with the majority of drugs localized near the origin, indicating no activity affecting either condition. The red diagonal identity line indicates where drugs affect PC9 cells equally under both treatment conditions. The majority of drugs have no significant effect under either condition. The color-coded density gradient represents 95% of drugs with outliers shown as blue dots. The indicated 95% tolerance interval (TI) was constructed after fitting the joint density to a single two-dimensional Gaussian distribution. Drugs indicated in blue or red were identified as hits based on statistical significance and >2-fold change in effect size between the two treatment conditions (see Fig. 1C). Drugs highlighted in blue (erlotinib, gefitinib) significantly enhanced macrophage-dependent killing of GFP+ PC9 cells. Drugs highlighted in red resulted in more cancer cell growth of PC9 cells in the presence of macrophages+anti-CD47 therapy versus the PC9 alone condition. However, drugs within this category exhibited different effects. Anthracyclines (**B**) and other chemotherapy drugs (**C**) exerted direct cytotoxicity to the PC9 cells alone, but the addition of macrophages+anti-CD47 protected the cancer cells from these drugs. In contrast, steroids (**D**) and retinoids (**E**) inhibited macrophage-mediated killing of the PC9 cells, thereby resulting in relatively enhanced growth in the macrophage+anti-CD47 condition. (B-E) Highlighted classes of drugs are show in red.

**Supplemental Figure 4:**
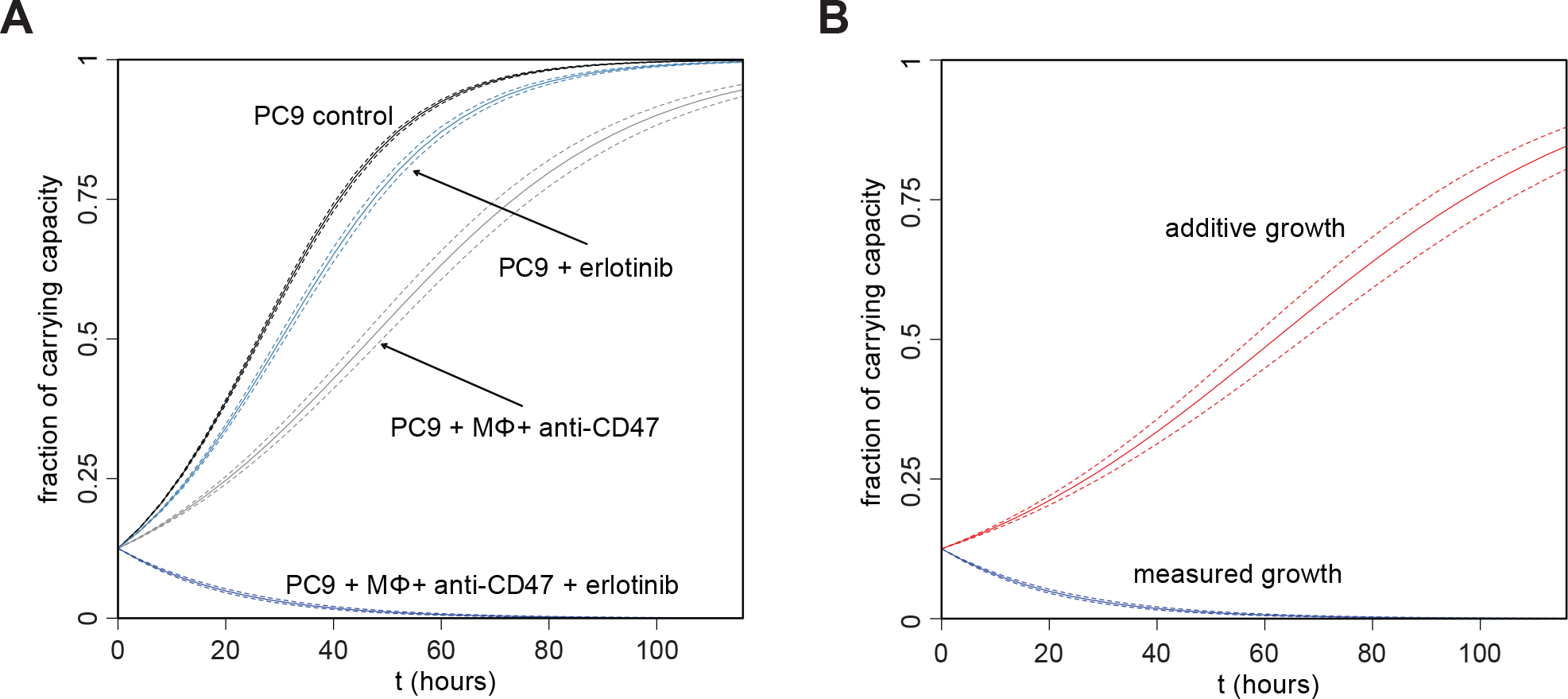
Modeling additive growth reveals synergy between EGFR inhibitors and anti-CD47 therapy. (**A**) Growth rates were estimated by fitting logistic growth models to time series data collected on PC9 cells grown under various conditions. Because the presence of macrophages with anti-CD47 antibodies elicits an anti-tumor response that limits the carrying capacity of PC9 cultures, the plots for fitted growth are rendered in terms of cell populations (i.e. area of GFP+ cells) normalized to carrying capacity. (**B**) The estimated growth rates were used to construct a purely additive model (i.e. without any synergy or antagonism) for the combined effects of erlotinib and macrophages + anti-CD47 antibodies. The dashed lines for each curve indicate the standard errors in the fitting. The dramatic reduction observed in the growth rate of PC9 cells due to the combination of erlotinib with macrophages + anti-CD47 antibodies is far greater than that predicted under the assumption of additivity, indicating that cooperativity is present.

**Supplemental Figure 5:**
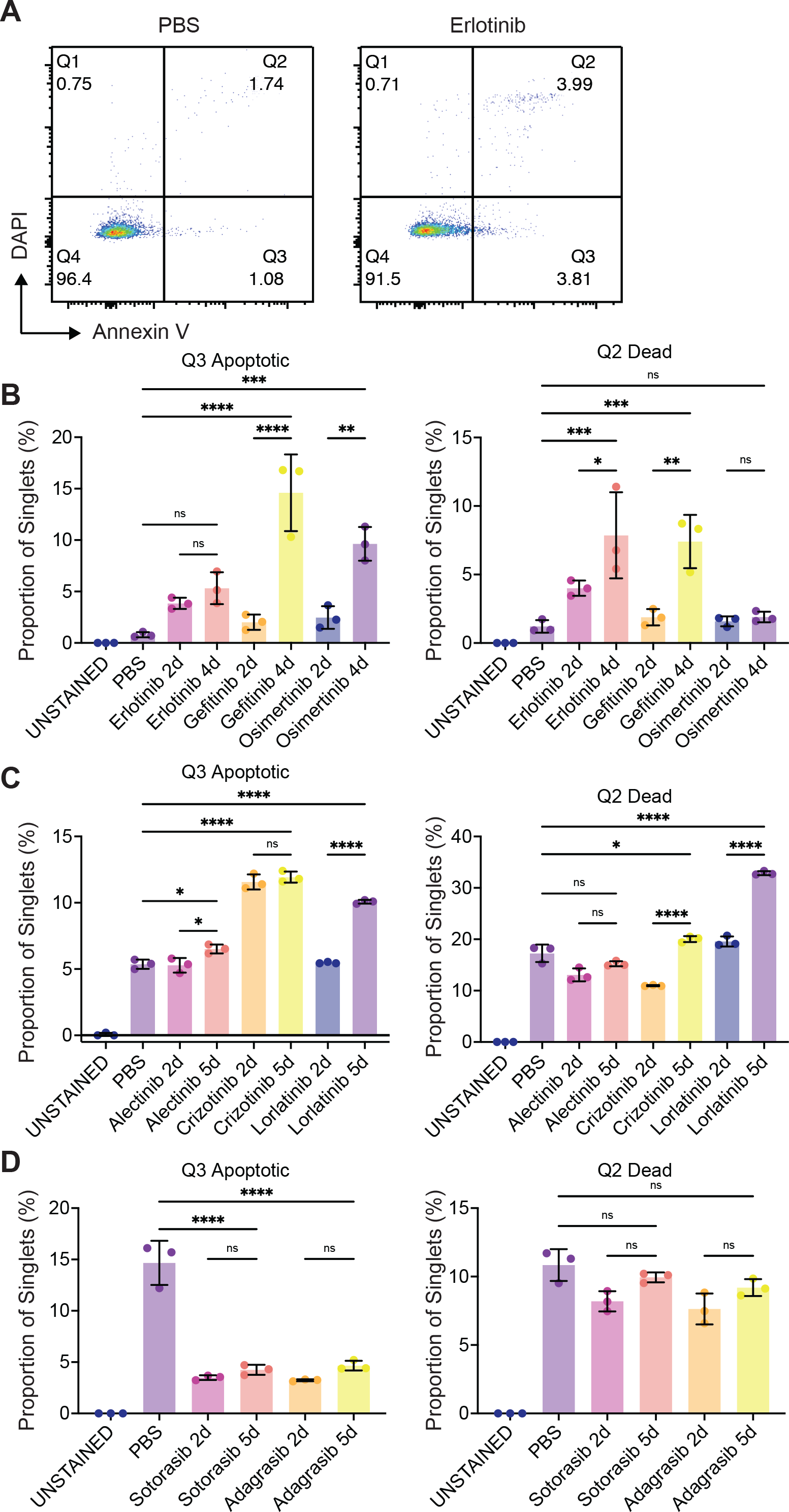
Analysis of apoptosis and cell death in response to targeted therapies. Lung cancer cells were treated with the indicated targeted therapies for 2-5 days. Adherent cells were collected and analyzed by flow cytometry for viability and apoptosis using annexin V and DAPI. (**A**) Representative plots showing PC9 cells treated with vehicle control or erlotinib to demonstrate gating strategy. Gates were drawn to quantify apoptotic cells (annexin V+, DAPI-) and dead cells (annexin V+, DAPI+). (**B**) Quantification of the percentage of PC9 cells undergoing apoptosis or cell death in response to the indicated EGFR TKIs. (**C**) Quantification of the percentage of NCI-H3122 cells undergoing apoptosis or cell death in response to the indicated ALK TKIs. (**D**) Quantification of the percentage of NCI-H358 cells undergoing apoptosis or cell death in response to the indicated KRAS^G12C^ inhibitors. (B-D) Data represent mean ± SD from 3 replicates performed from one experiment. ns, not significant, *p<0.05, **p<0.01, ***p<0.001, ****p<0.0001 by one-way ANOVA with Tukey’s multiple comparisons test.

**Supplemental Figure 6:**
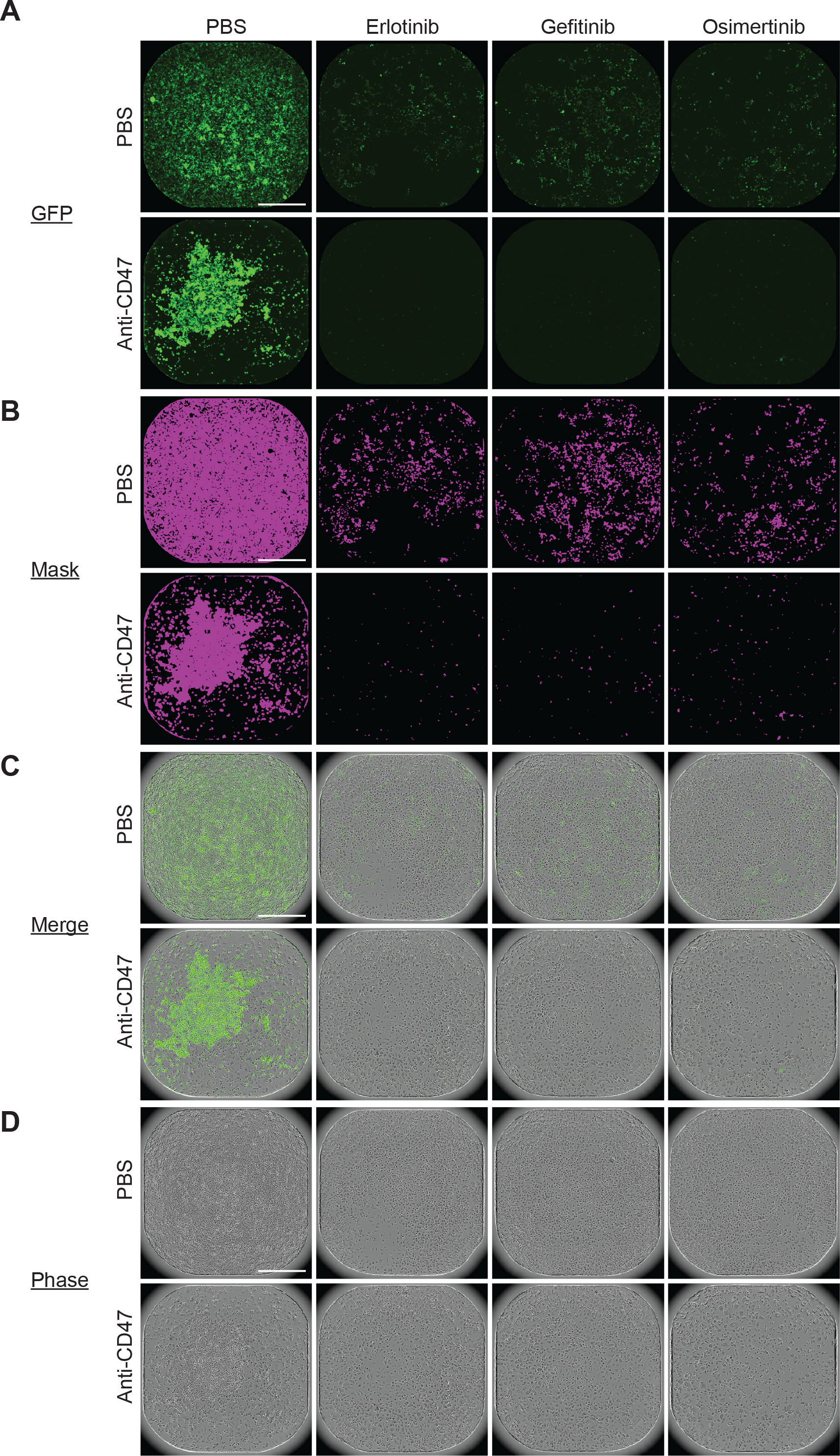
Representative images of long-term co-culture assays using GFP+ PC9 cells and human macrophages. GFP+ PC9 cells were combined with primary human macrophages and the indicated drug therapies in 384-well plates. Representative images are shown from a macrophages derived from an individual blood donor at t = 6d 12h. **(A)** Whole well imaging of the GFP+ channel from wells treated with the indicated therapies. (**B**) Image mask of GFP+ pixels (purple) used for quantification and analysis. (**C**) Overlay of GFP+ channel with phase contrast imaging. (**D**) Phase contrast imaging showing confluency of wells with GFP+ PC9 cells and primary human macrophages present.

**Supplemental Figure 7:**
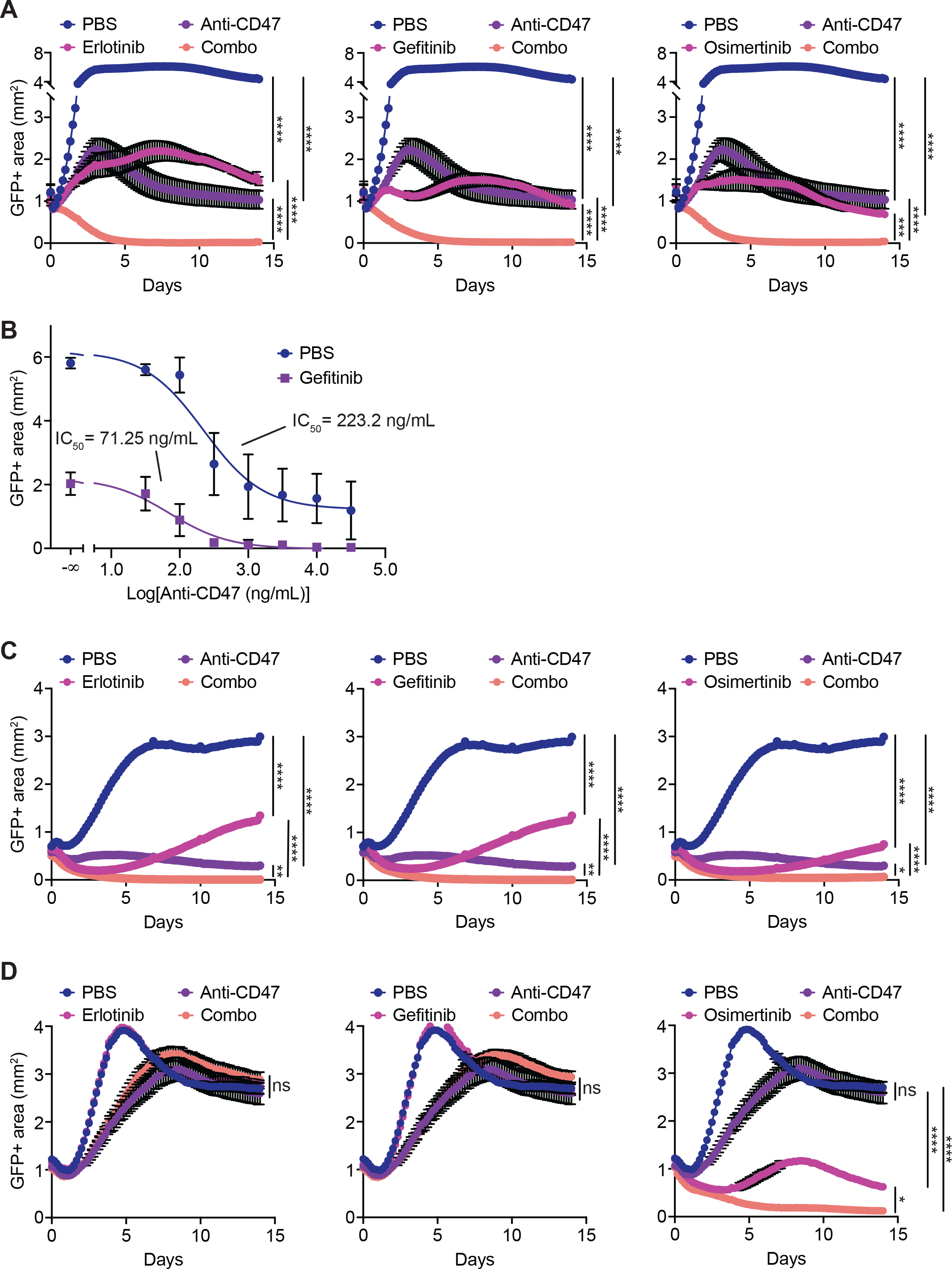
Growth curves of long-term assays using human macrophages and different *EGFR* mutant lung cancer specimens. GFP+ lung cancer cells were combined with primary human macrophages and the indicated drug therapies in 384-well plates. The GFP+ area, representing the growth or death of the GFP+ cancer cells, was evaluated by whole-well imaging every 4 hours and quantified by automated image analysis. (**A**) GFP+ PC9 cells co-cultured with macrophages and erlotinib (left), gefitinib (middle), or osimertinib (right). **(A)** Co-culture assays using GFP+ PC9 cells and human macrophages to evaluate a dose-response relationship. The concentration of anti-CD47 was titrated alone or in combination with gefitinib at 100 nM. The IC_50_ for anti-CD47 improved from 223.2 ng/mL (95% CI 158.2-317.3) to 71.25 ng/mL (95% CI 52.39-97.22). GFP+ area measured and compared on day 6.5 of co-culture. (**C**) GFP+ MGH119 cells co-cultured with macrophages and erlotinib (left), gefitinib (middle), or osimertinib (right). (**D**) GFP+ MGH134 cells co-cultured with macrophages and erlotinib (left), gefitinib (middle), or osimertinib (right). (A-D) Data at each timepoint represent mean ± SEM from 3-4 technical replicates per donor and n = 4-8 independent macrophage donors. ns, not significant, *p<0.05, **p<0.01, ***p<0. 001, ****p<0.0001 by two-way ANOVA with Holm-Sidak multiple comparisons test on day 14 of co-culture or as indicated.

**Supplemental Figure 8:**
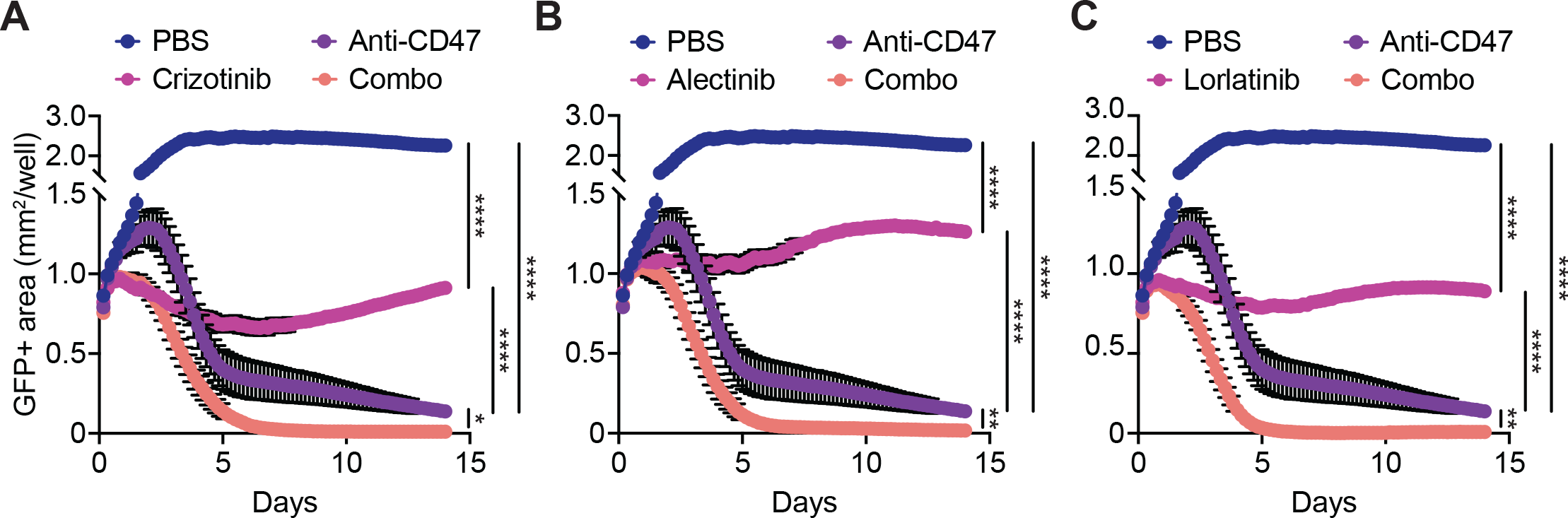
Growth curves of long-term assays using human macrophages and an *ALK* rearranged lung cancer cell line. GFP+ NCI-H3122 lung cancer cells were combined with primary human macrophages and the indicated drug therapies in 384-well plates. The GFP+ area, was evaluated by whole-well imaging every 4 hours and quantified by automated image analysis. GFP+ NCI-H3122 cells co-cultured with macrophages and crizotinib (**A**), alectinib (**B**), or lorlatinib (**C**). Data at each timepoint represent mean ± SEM from 3 technical replicates per donor using n = 4 independent macrophage donors. ****p<0.0001 by two-way ANOVA with Holm-Sidak multiple comparisons test on day 14 of co-culture.

**Supplemental Figure 9:**
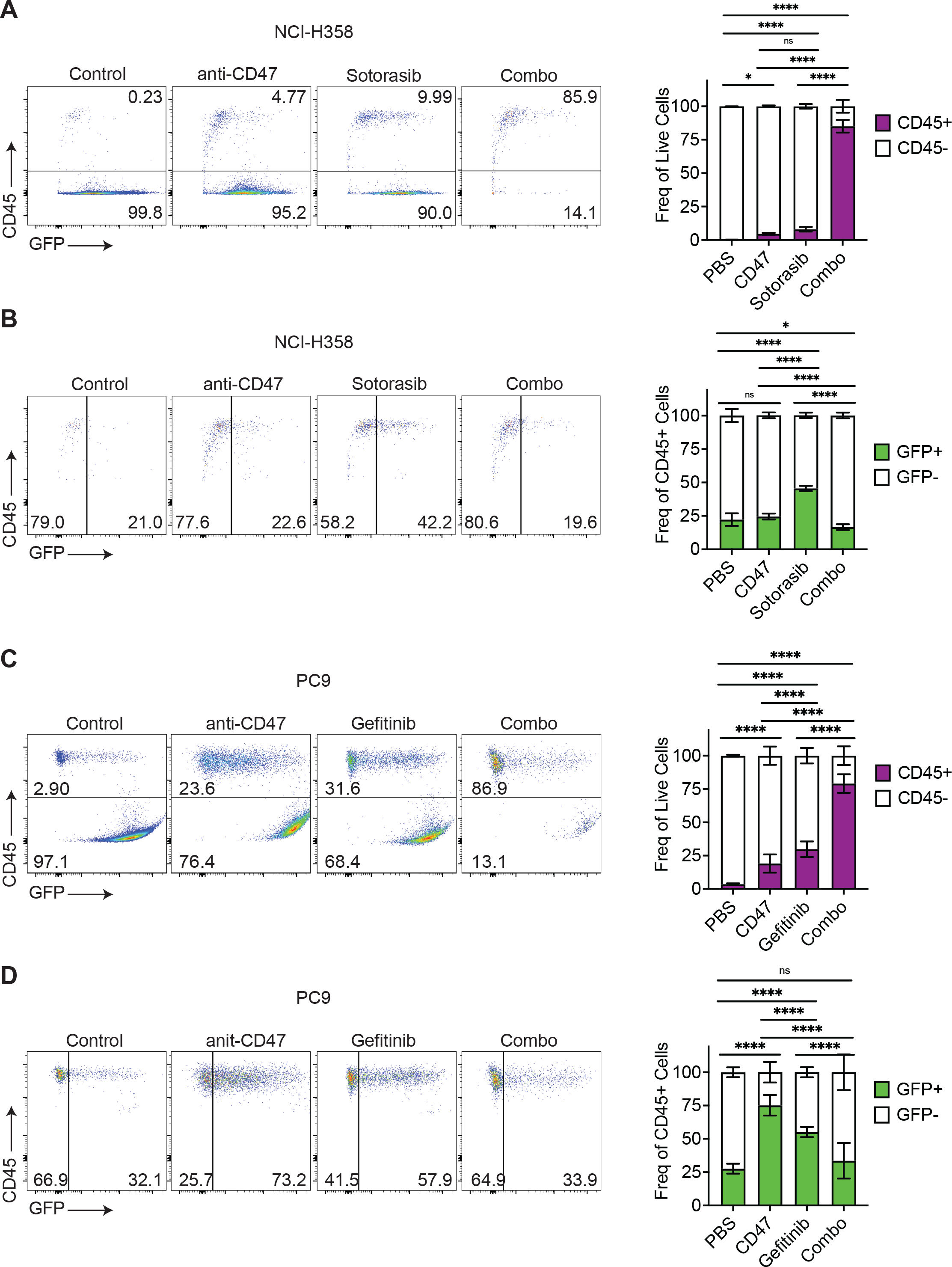
Flow cytometry analysis of long-term co-cultures assays demonstrates phagocytosis and elimination of cancer cells. Primary human macrophages were co-cultured with GFP+ NCI-H358 cells (**A-B**) or GFP+ PC9 cells (**C-D**) and the indicated therapies. Cells were collected on day 4 of co-culture and analyzed by flow cytometry. Macrophages were identified by APC anti-CD45 and lung cancer cells were identified by GFP fluorescence. The percentage of GFP+ macrophages was quantified as a representation of phagocytosis. An increase in the percentage of CD45+ cells (A, C) was observed due to elimination of cancer cells in the co-culture. Similarly, the percentage of phagocytic macrophages decreased in the combo therapy treatment due to decreases in cancer cell number and digestion of internalized material (B, D). For each condition, representative FACS plots are shown on the left and summary bar graphs depicting mean ± SD are shown on the right. ns, not significant, *p<0.05, ****p<0.0001 by two-way ANOVA with Tukey’s multiple comparisons test from one experiment using macrophages derived from 3 independent donors with 8 technical replicates per donor (PC9) or 1 donor with 8 technical replicates (NCI-H358).

**Supplemental Figure 10:**
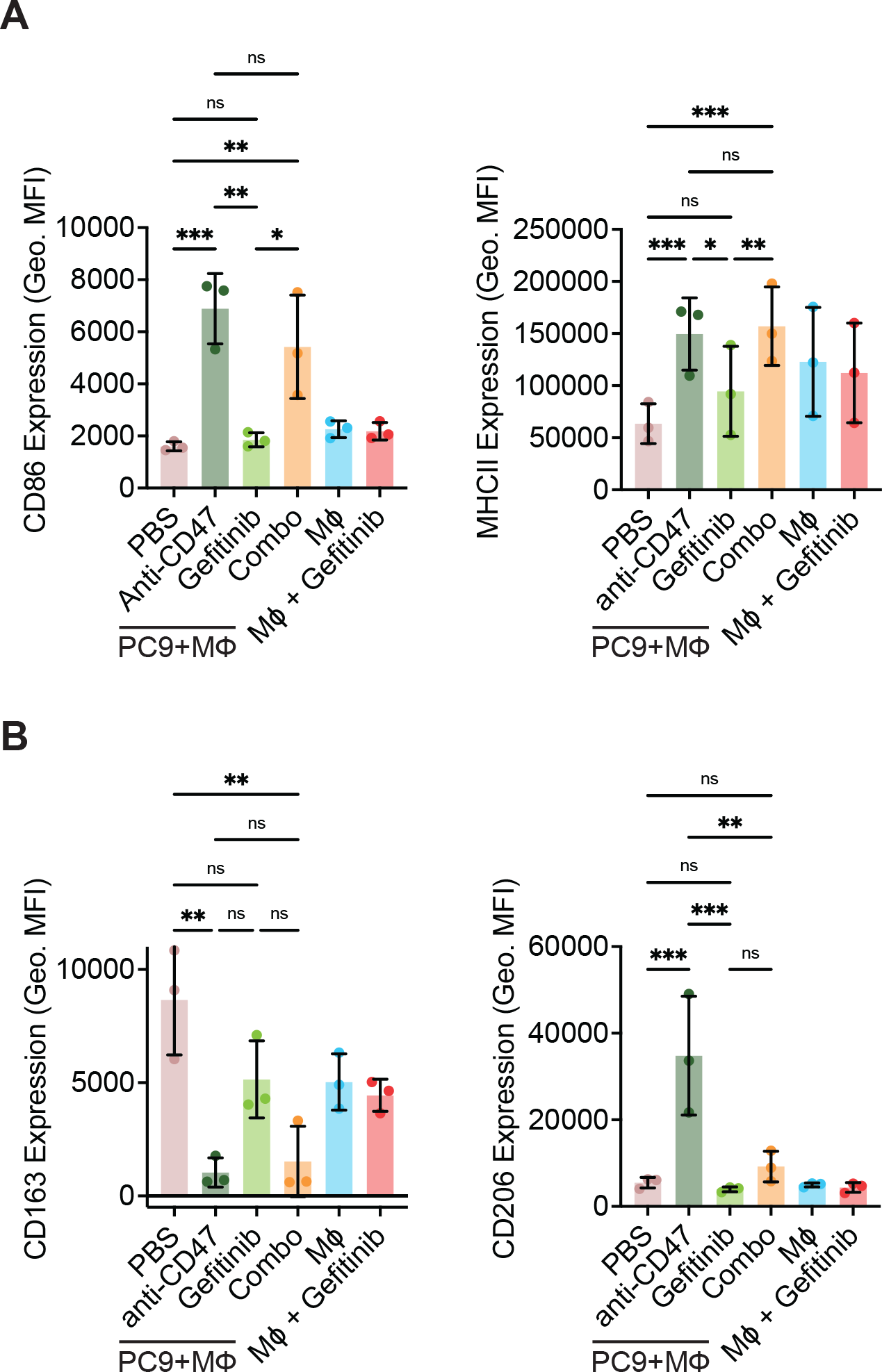
Analysis of macrophage polarization state following co-culture of macrophages and lung cancer cells. Primary human macrophages were cultured alone or with GFP+ PC9 cells and the indicated therapies. Cells were collected on day 4 of co-culture and analyzed by flow cytometry for the antigens associated with (**A**) M1 polarization (CD86, MHC II); or (**B**) M2 polarization (CD163, CD206). Bar graphs depict mean ± SD from n = 3 independent donors with one technical replicate per donor. ns, not significant, *p<0.01, ****p<0.0001 by two-way ANOVA with Tukey’s multiple comparisons test from one independent experiment.

**Supplemental Figure 11:**
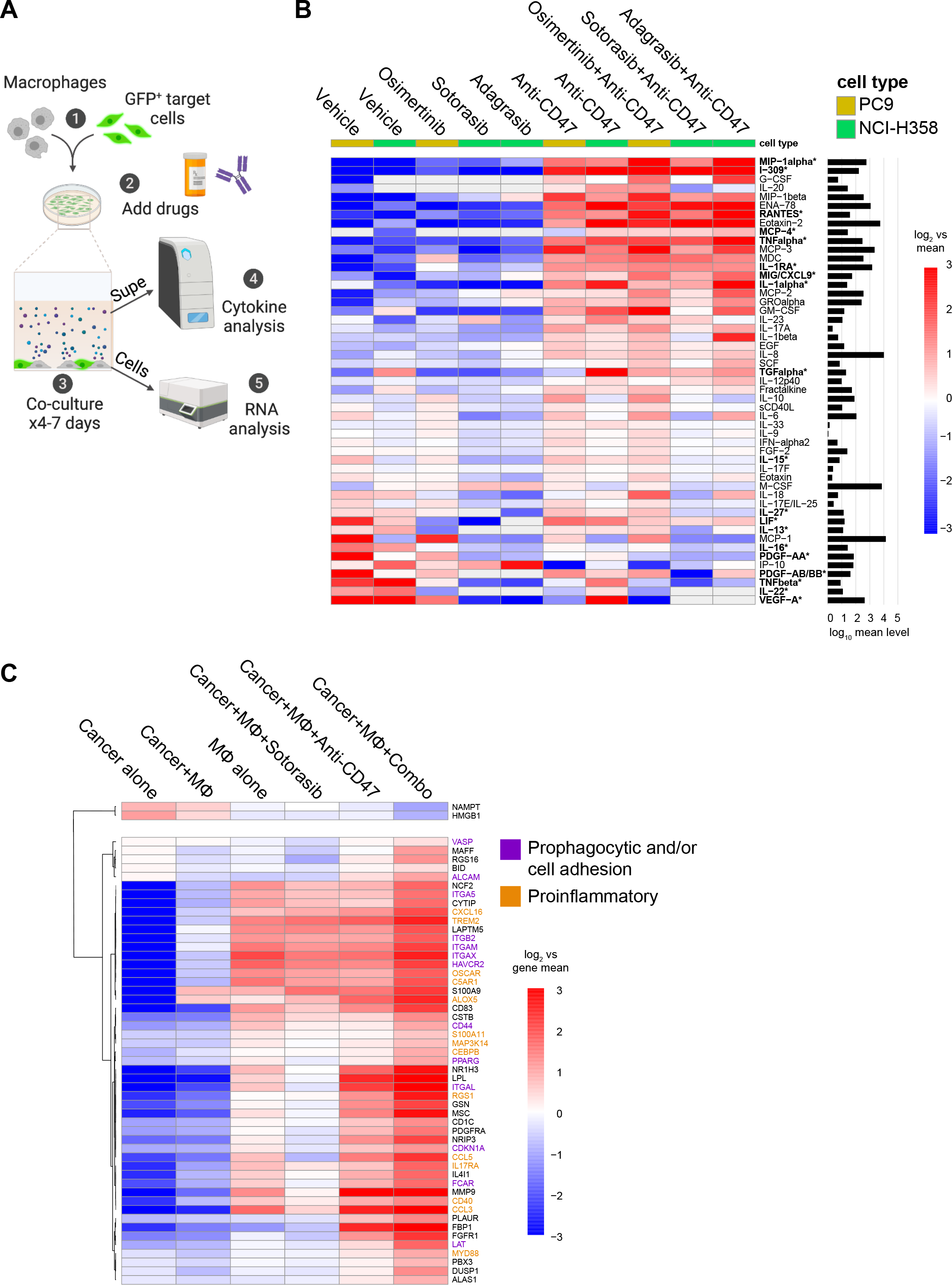
The combination of targeted therapies and anti-CD47 elicits unique cytokine and gene expression signatures in co-culture assays. (**A**) Diagram showing experimental setup of cytokine and RNA profiling experiments. Primary human macrophages were co-cultured with GFP+ target NSCLC cells (PC9 or NCI-H358) with targeted therapies and/or an anti-CD47 antibody. Cells were co-cultured for 4-7 days. Supernatants were collected and subjected to multiplex cytokine analysis of 71 human analytes by addressable laser bead immunoassay. In a separate experiment, adherent cells were collected and subjected to targeted gene expression profiling of 770 myeloid-derived genes using an nCounter Myeloid Innate Immunity Panel (Nanostring). (**B**) Multiplex cytokine analysis of supernatants from co-culture assays using primary human macrophages and PC9 or NCI-H358 NSCLC cells. Each column represents data from the indicated drug treatments, and cytokines levels were compared by mean fluorescence intensity. Cytokines that were statistically significant by ANOVA (FDR <0.05) across experiments between anti-CD47 therapy and combo therapy groups are indicated in bold with an asterisk. Color scale indicates log2 fold-change versus mean for each individual cytokine, and the mean level for each cytokine is shown in the bar graph on the right. Data represent specimens collected on day 4 and day 7, with each time point containing 4 technical replicates from three independent donors mixed in equal ratios (PC9 experiment), or 1 technical replicate from each of 4 independent donors tested individually or mixed in equal ratios. (**C**) Targeted gene expression analysis depicting myeloid-derived genes from co-culture assays of primary human macrophages and NCI-H358 cells. Heatmap indicates hierarchical clustering of genes that were significantly downregulated or upregulated following treatment with the combination of sotorasib and anti-CD47 therapy versus all other treatment groups by ANOVA (FDR <0.05). Color scale indicates log2 fold-change versus mean for each individual gene. Data represent analysis performed with 4 independent donors with one technical replicate per donor with specimens collected on day 4 of co-culture. Genes associated with phagocytosis and/or cell-cell adhesion are depicted in purple. Genes that have reported proinflammatory functions are depicted in orange.

**Supplemental Figure 12:**
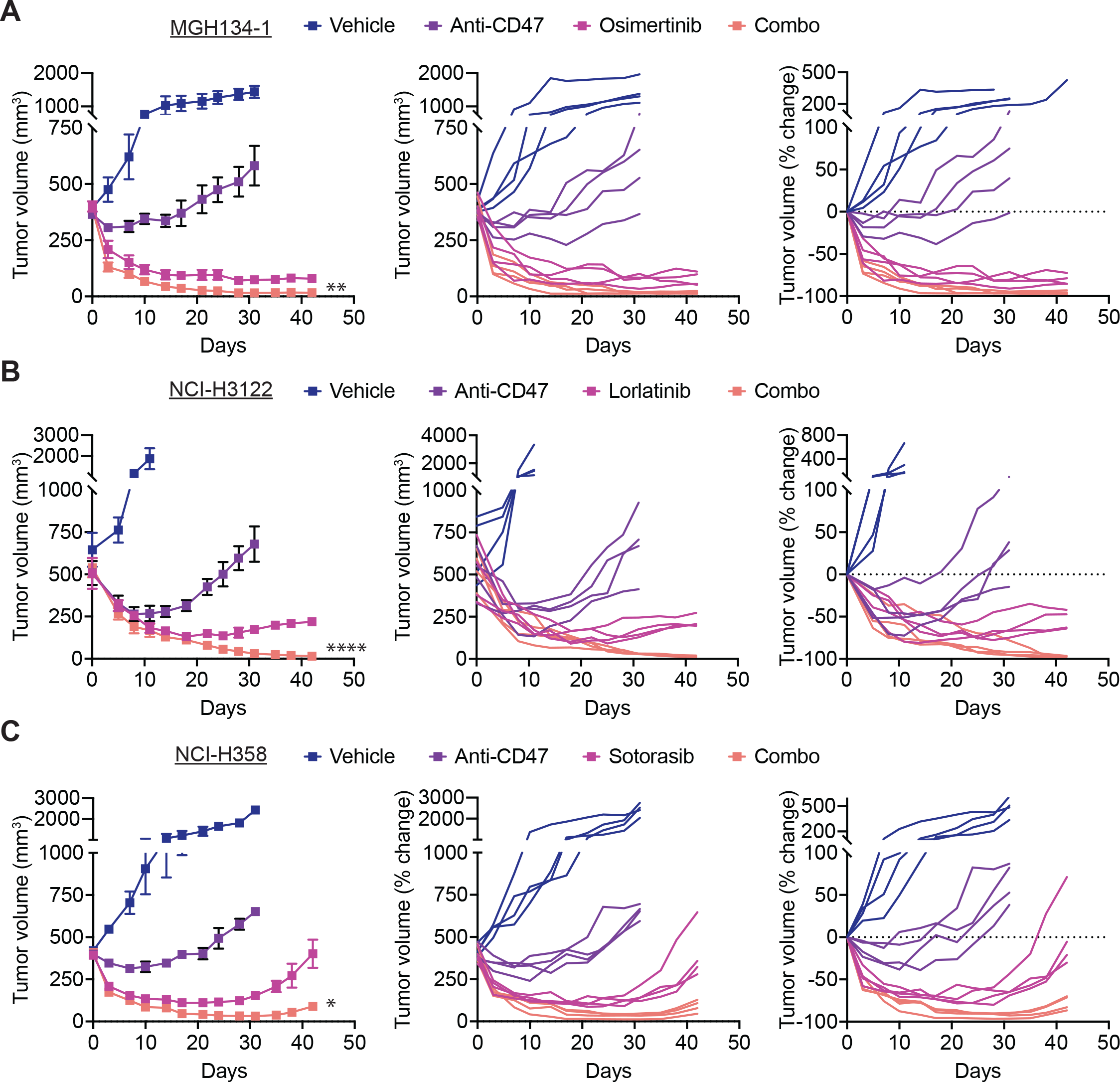
Growth curves of lung cancer tumors in xenograft treatment experiments. Full growth curves from mouse xenograft tumor models shown in Fig. 5 depicting tumor volumes as mean ± SEM (left), growth curves from individual mice (middle) or fold-change from individual mice. (**A**) GFP+ MGH134-1 cells treated with vehicle control, anti-CD47 alone, osimertinib alone, or the combination (combo). (**B**) GFP+ NCI-H3122 cells treated with vehicle control, anti-CD47 alone, lorlatinib alone, or the combination. (**C**) GFP+ NCI-H358 cells treated with vehicle control, anti-CD47 alone, sotorasib alone, or the combination. (A-C) *p<0.05, **p<0.01, ***p<0.001 for the combination therapy versus targeted therapy by unpaired t test. For each experiment, n = 4 mice per treatment cohort.

**Supplemental Figure 13:**
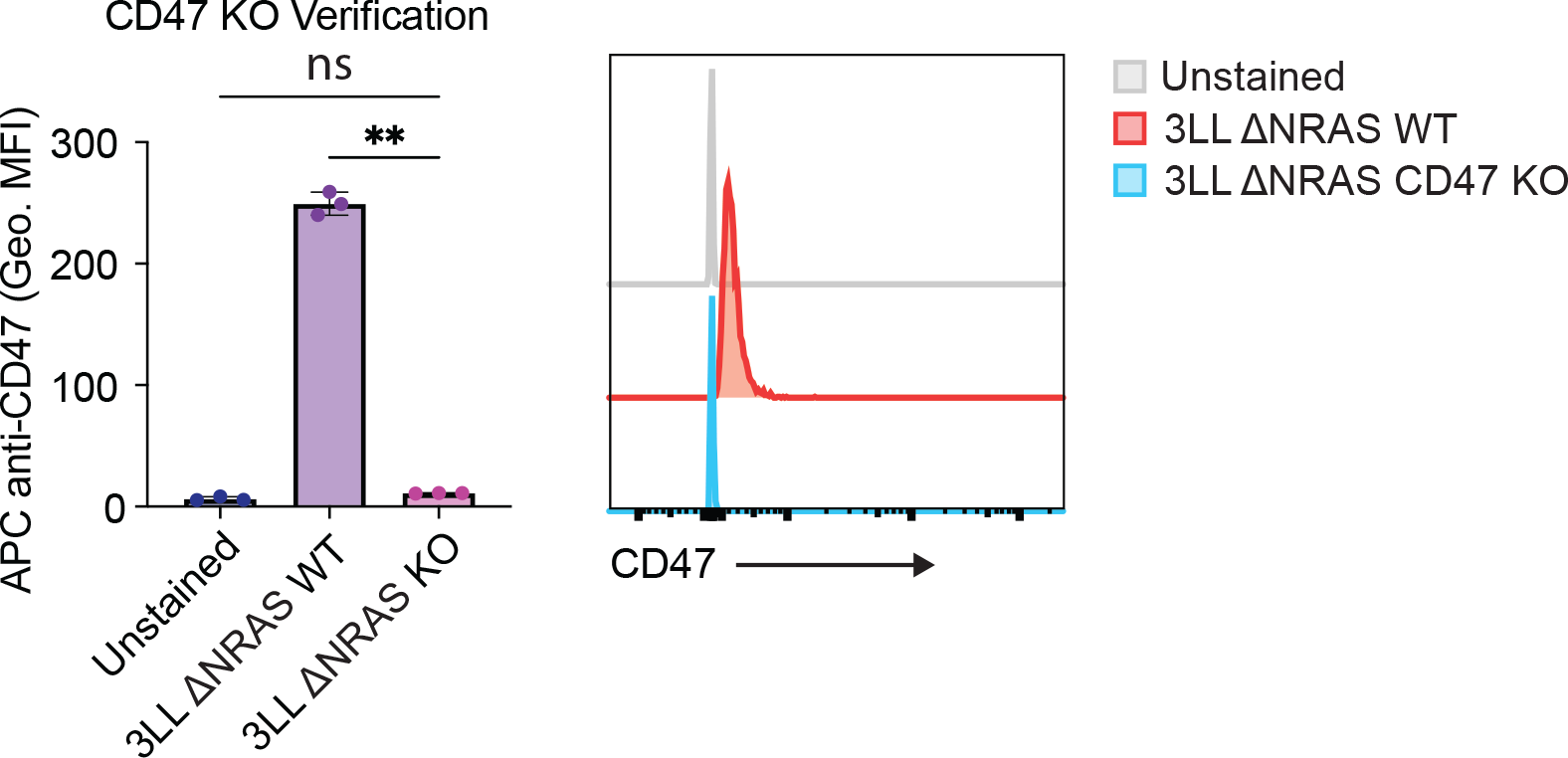
Validation of a CD47 KO line generated by CRISPR/Cas9 editing of 3LL ΔNRAS cells. Cell-surface CD47 expression as detected by flow cytometry for wild-type (WT) 3LL ΔNRAS cells or a CD47 knockout (KO) variant. Data are depicted as mean ± SD from three replicates (left), or as representative histograms (right). ns, not significant, **p<0.01 by one-way ANOVA with Holm-Sidak multiple comparisons test.

**Supplemental Figure 14:**
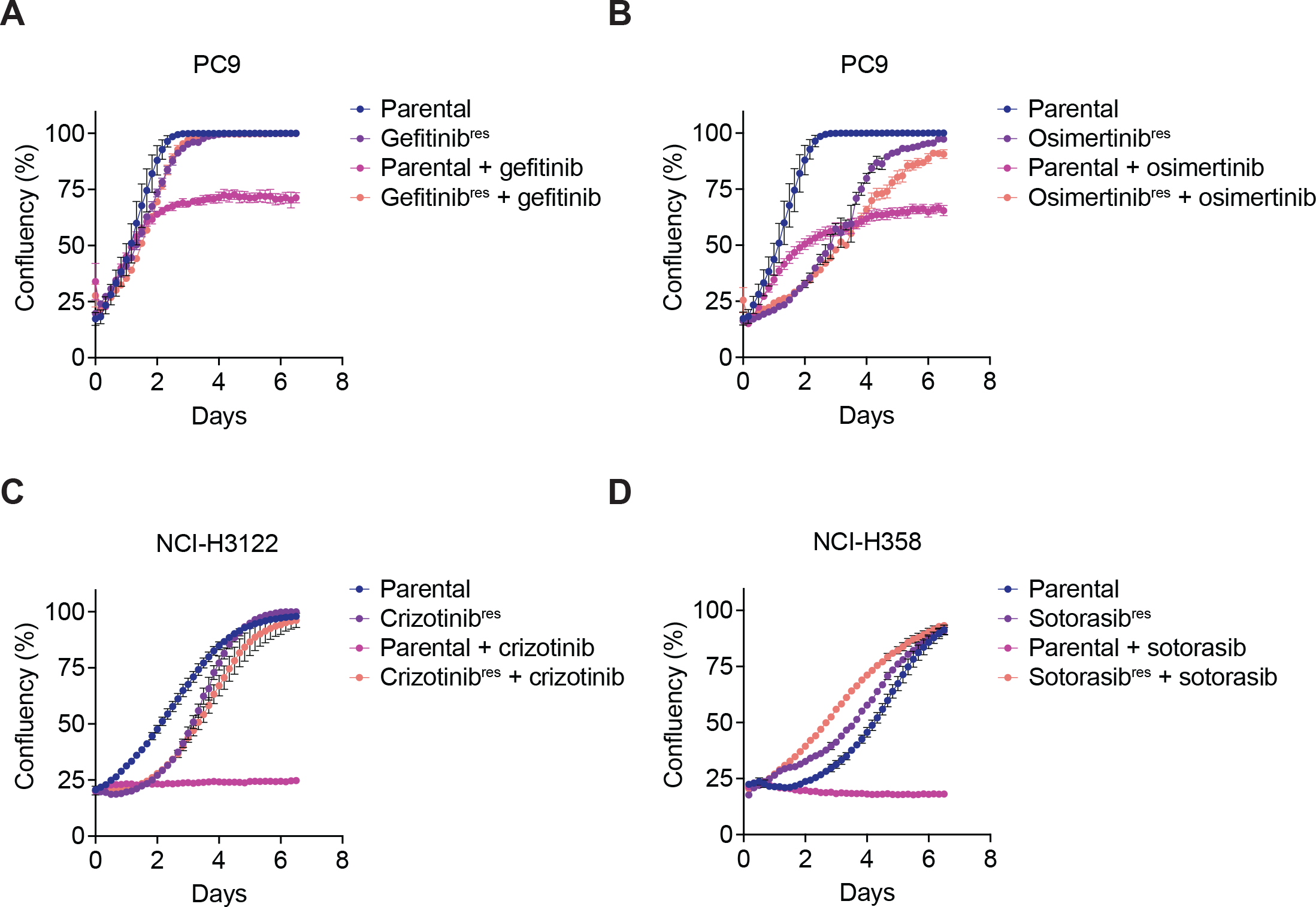
Proliferation of NSCLC cell lines in vitro after acquiring resistance to targeted therapies. Resistant cell lines were generated by prolonged culture of NSCLC cell lines in appropriate targeted therapy. Proliferation was evaluated by confluency analysis as measured by phase microscopy and automated image analysis. Proliferation was measured without drug selection or with 1 uM targeted therapy as indicated. Cell lines tested included PC9 cells resistant to gefitinib (**A**) or osimertinib (**B**), NCI-H3122 cells resistant to crizotinib (**C**), or NCI-H358 cells resistant to sotorasib (**D**). For the majority of cell lines, growth rates were comparable between parental and resistant cells in the absence of targeted therapy and approached 100% confluency by day 6.5 of culture. Data represent mean of 3 technical replicates ± SEM from one independent experiment for each cell line. (A-B) PC9 evaluation was performed in a single experiment and separated into distinct plots with the same parental curve reproduced for data visualization.

**Supplemental Figure 15:**
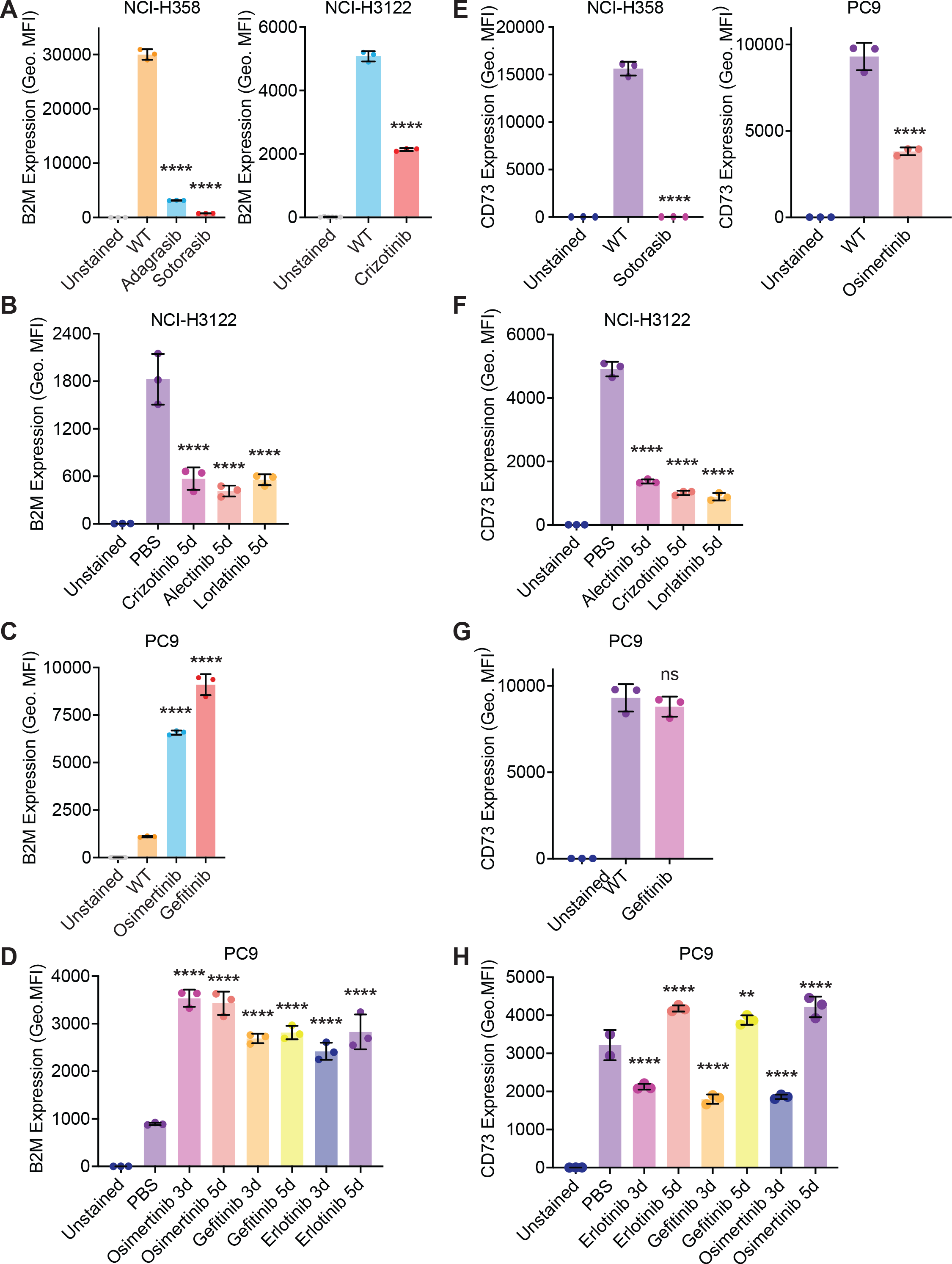
Changes in B2M and CD73 expression on lung cancer cells exposed to targeted therapies. (**A**) Downregulation of B2M on NCI-H358 cells or NCI-H3122 cells resistant to the indicated targeted therapies. (**B**) Downregulation of B2M on NCI-H3122 cells following treatment with the indicated ALK inhibitors. (**C**) B2M was not downregulated on PC9 cells that are resistant to EGFR inhibitors, nor PC9 cells exposed to EGFR inhibitors in culture (**D**). (**E**) CD73 is downregulated on NCI-H358 and PC9 cells that are resistant to the indicated targeted therapies. (**F**) NCI-H3122 cells downregulate CD73 in response to the indicated targeted therapies. (**G**) PC9 cells resistant to gefitinib did not downregulate CD73. (**H**) CD73 is dynamically regulated on the surface of PC9 cells in response to EGFR inhibitors, with initial downregulation after 3 days of exposure, followed by increased surface expression. (A-H) Data represent mean ± SD from 3 technical replicates from individual experiments. ns, not significant, ****p<0.0001 by one-way ANOVA with Holm-Sidak multiple comparisons test.

**Supplemental Figure 16:**
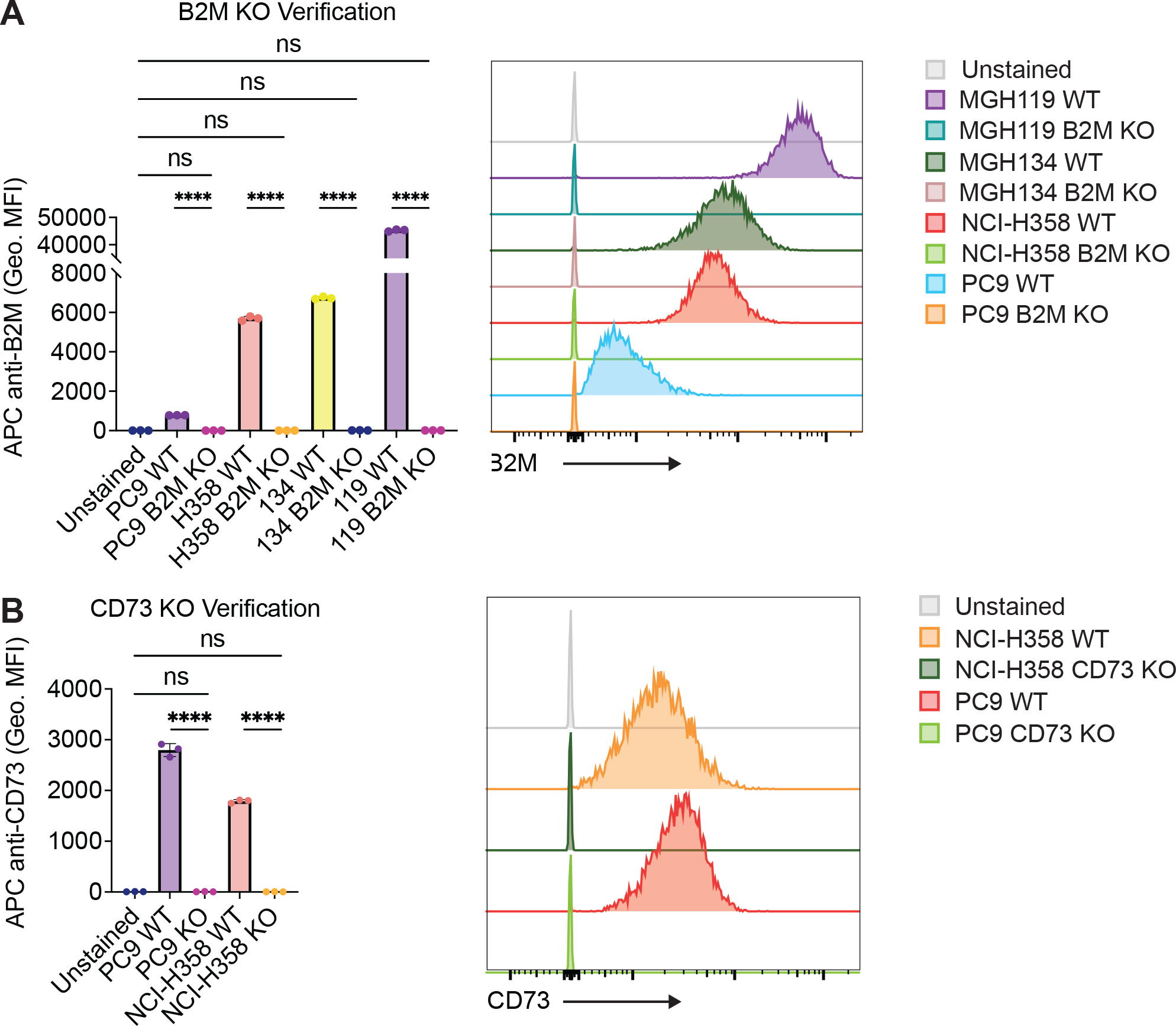
Validation of B2M KO and CD73 KO lines generated by CRISPR/Cas9 editing of human lung cancer cell lines. (**A**) Flow cytometry analysis of B2M expression on the surface of wild-type (WT) PC9, NCI-H358, MGH134, and MGH119 cells compared to their respective B2M KO variants. Left, quantification of geometric mean fluorescence intensity (Geo. MFI). Right, representative histograms showing B2M surface expression. (**B**) Flow cytometry analysis of CD73 expression on the surface of wild-type (WT) PC9 and NCI-H358 cells compared to their respective CD73 KO variants. Left, quantification of geometric mean fluorescence intensity (Geo. MFI). Right, representative histograms showing B2M surface expression. (A-B) Data represent mean ± SD from 3 technical replicates from one individual experiment. ns, not significant, ****p<0.0001 by one-way ANOVA with Holm-Sidak multiple comparisons test.

**Supplemental Figure 17:**
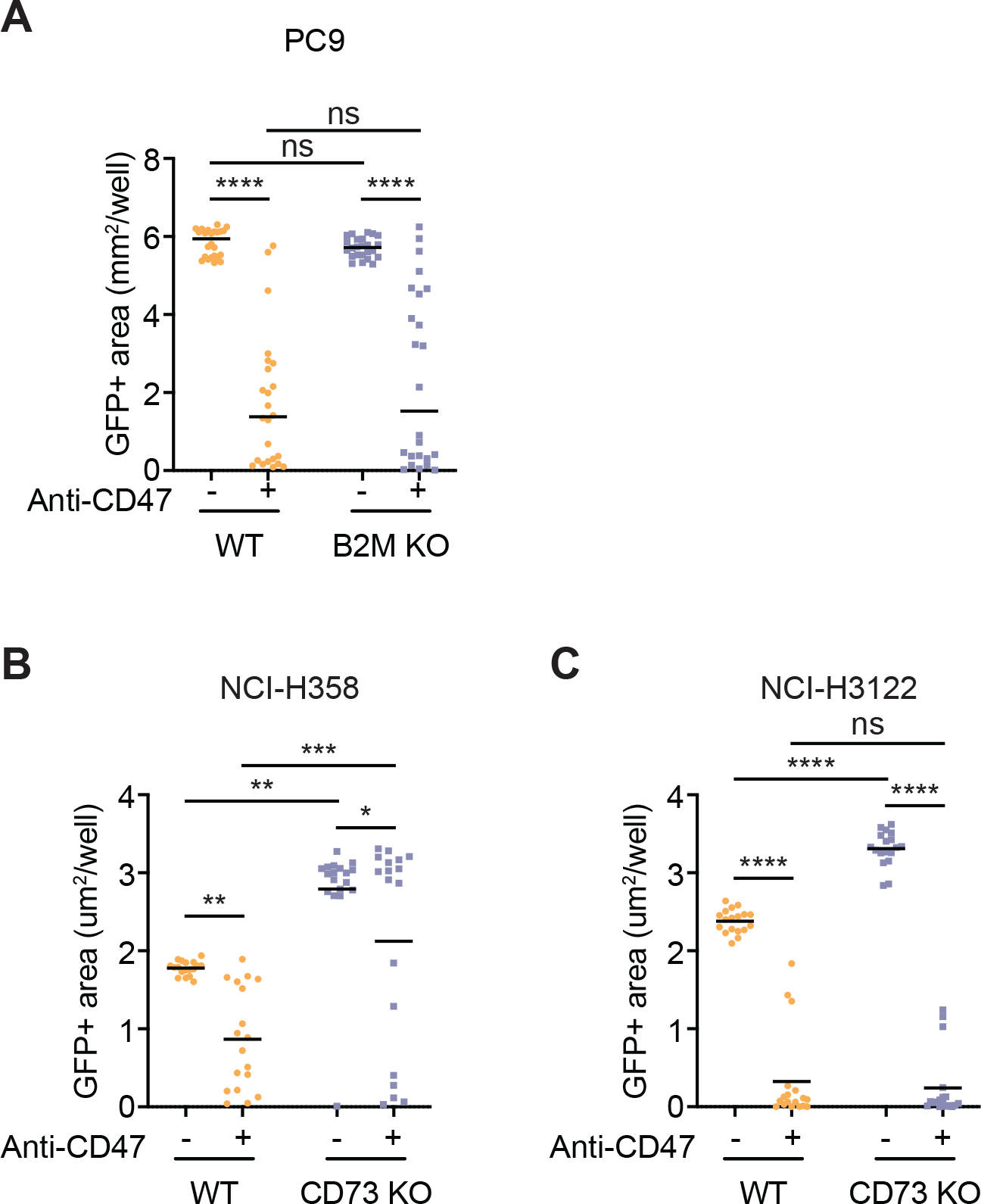
Genetic deletion of B2M or CD73 does not make some NSCLC cell lines more vulnerable to macrophage attack. (**A**) Evaluation of wild-type versus B2M KO PC9 cells in long-term co-culture assays with human macrophages. Cells were treated with vehicle control or an anti-CD47 antibody. Data are combined from two independent experiments performed with a total of n = 8 independent macrophage donors with 3 technical replicates per donor. (**B-C**) Evaluation of wild-type versus CD73 KO NCI-H358 (B) or NCI-H3122 cells (C) in long-term co-culture assays with human macrophages. Cells were treated with vehicle control or an anti-CD47 antibody. Data represent mean ± SD from n = 6 independent macrophage donors with 3 technical replicates per donor from one experiment. (A-C) ns, not significant, *p<0.05, **p<0.01, ***p<0.001, ****p<0.0001 by two-way ANOVA with Holm-Sidak multiple comparisons test.

**Supplemental Movie 1: Long-term co-culture assays using GFP+ PC9 cells and human macrophages treated with vehicle control.** Whole-well imaging was performed of GFP+ channel showing growth of *EGFR* mutant cancer cells in the presence of primary human macrophages (unlabeled). GFP+ cancer cells growth without any substantial inhibition by macrophages. Video depicts 8 days of elapsed time. Images acquired every 4 hours. Scale bar = 1.0 mm.

**Supplemental Movie 2: Long-term co-culture assays using GFP+ PC9 cells and human macrophages treated with gefitinib as a single agent.** Whole-well imaging was performed of GFP+ channel showing growth of *EGFR* mutant cancer cells in the presence of primary human macrophages (unlabeled). GFP+ cancer cells are restricted in their growth, but persister cells always form and remain in the culture over time. Video depicts 8 days of elapsed time. Images acquired every 4 hours. Scale bar = 1.0 mm.

**Supplemental Movie 3: Long-term co-culture assays using GFP+ PC9 cells and human macrophages treated with anti-CD47 as a single agent.** Whole-well imaging was performed of GFP+ channel showing growth of *EGFR* mutant cancer cells in the presence of primary human macrophages (unlabeled). GFP+ cancer cells are attacked by macrophages, but foci of cancer cells remain in the culture over time. Video depicts 8 days of elapsed time. Images acquired every 4 hours. Scale bar = 1.0 mm.

**Supplemental Movie 4: Long-term co-culture assays using GFP+ PC9 cells and human macrophages treated with the combination of gefitinib and anti-CD47.** Whole-well imaging was performed of GFP+ channel showing growth of *EGFR* mutant cancer cells in the presence of primary human macrophages (unlabeled). GFP+ cancer cells are attacked and fully eliminated by macrophages such that persister cells do not form. Video depicts 8 days of elapsed time. Images acquired every 4 hours. Scale bar = 1.0 mm.

